# Deciphering the RNA Modification Landscape in Arabidopsis Chloroplast tRNAs and rRNAs Reveals a Blend of Ancestral and Acquired Characteristics

**DOI:** 10.1101/2024.06.14.598963

**Authors:** Kinga Gołębiewska, Pavlína Gregorová, L. Peter Sarin, Piotr Gawroński

## Abstract

Chloroplasts in plant leaves are essential for protein synthesis, relying on transfer RNAs (tRNAs) and ribosomal RNAs (rRNAs) encoded by the chloroplast genome. Although post-transcriptional modifications of these non-coding RNAs are common in many systems, chloroplast tRNA and rRNA modifications are not well characterised.

In this study, we investigated the post-transcriptional modifications in chloroplast tRNAs and rRNAs of *Arabidopsis thaliana* using tRNA sequencing, liquid chromatography-mass spectrometry, targeted rRNA sequencing, and analysis of public data.

Our results revealed similarities between chloroplast non-coding RNAs and bacterial systems (e.g., *Escherichia coli*), such as modification patterns at the anticodon-adjacent position and the variable loop of tRNAs, along with conserved modifications in the small subunit rRNA. Additionally, we identified features shared with eukaryotic systems that likely contribute to the correct three-dimensional structure of chloroplast tRNAs. Unique modifications were also discovered, including a potential novel modification at wobble position in tRNA-Ile^CAU^, which may be crucial for distinguishing isoleucine codons from methionine codons, and chloroplast-specific rRNA modifications that likely compensate for altered ribosome structure.

These findings suggest that the chloroplast translation machinery, through co-evolution with its eukaryotic host, has adopted features beyond those typically found in bacteria, reflecting a blend of ancestral and acquired characteristics.

## INTRODUCTION

Chloroplasts, found in plant and algal cells, are organelles derived from ancient cyanobacteria (1). These semi-independent organelles contain their own genetic material, known as plastome, which is co-expressed with the nuclear genome. The plastome of land plants typically spans from 120 to 160 kb and encodes approximately 120 genes. Many of these plastid genes are vital as they encode components of the photosynthetic apparatus and gene expression machinery, such as RNA polymerase subunits, ribosomal proteins, ribosomal RNAs (rRNAs) and transfer RNAs (tRNAs). The plastome encodes all RNAs necessary for translation within chloroplasts, as importing these molecules into chloroplasts is unlikely. Interestingly, unlike their prokaryotic counterparts, regulation of chloroplast gene expression primarily occurs at post-transcriptional stages, particularly during translation (2). Recent research has provided a detailed understanding of the structure of chloroplast ribosomes (chloro-ribosomes), laying the groundwork for further characterisation of the chloroplast translation machinery and its regulation (3–6).

A chloroplast ribosome (70S) is a complex molecular assembly composed of rRNA and numerous proteins. It is composed of two primary subunits: the Small Subunit (SSU) – 30S, and the Large Subunit (LSU) – 50S. Each subunit contains distinct regions contributing to the ribosome’s overall functionality. The structural organisation of chloroplast ribosomes closely resembles that of prokaryotic ones, indicating a shared ancestry between chloroplasts and bacteria. However, subtle but significant variations exist between chloroplast and prokaryotic ribosomes. Notably, chloro-ribosomes feature unique rRNA sequences and protein compositions that differentiate them from their bacterial counterparts (3). These differences may confer distinct functional properties, adapting the ribosome to the specific chloroplast biology.

The SSU contains the 16S rRNA, housing the decoding center (DC). Furthermore, the subunit includes specific sites interacting with tRNAs. Notably, the A (aminoacyl) site accommodates incoming aminoacyl-tRNA during translation elongation, the P (peptidyl) site binds peptidyl-tRNA carrying the nascent polypeptide chain, and the E (exit) site assists deacyl-tRNA before dissociation from the ribosome. The primary function of SSU is to control the interaction between mRNA codons and tRNA anticodons, ensuring proper base-pairing and selection of the correct tRNA. The chloroplast LSU consists of 23S rRNA, 4.5S rRNA (a fragment derived from 3’ end of bacterial 23S), and 5S rRNA. This subunit also interacts with tRNAs, but it is primarily responsible for peptide bond formation between amino acids and facilitating the elongation of the polypeptide chain during protein synthesis.

The translation machinery within chloroplast includes rRNA and tRNAs that contain numerous post-transcriptionally modified nucleosides at key positions, a feature commonly conserved across living organisms. However, the functional significance of these RNA modifications varies considerably; the absence of some can lead to lethality, while the lack of others does not result in observable phenotype. This discrepancy is likely tied to the specific role of each modification. For instance, impaired modifications in the anticodon stem loop (ASL) of tRNAs can hinder translation efficiency, resulting in the aggregation and misfolding of essential proteins (7).

On the other hand, most rRNA modifications contribute to the appropriate tertiary structure of the molecule, typically enhancing protein synthesis efficiency without being strictly necessary for the process (reviewed in (8)). Moreover, RNA modifications can undergo stoichiometric changes in response to suboptimal environmental conditions (9, 10). The wide variety of RNA modifications is particularly prevalent in thermophilic organisms (11–13). Despite *Escherichia coli* being the most extensively studied prokaryotic organism in terms of RNA modifications, the functional implications of many of these modifications remain unclear (14). It is noteworthy that the set of RNA modifications present in bacteria varies depending on their adaptations to different environmental conditions.

tRNAs undergo extensive post-transcriptional modifications across all genetic systems. These modifications exhibit chemical diversity, with over 100 distinct modifications identified in tRNAs to date (14). Broadly, these modifications play critical roles in decoding codons in mRNA (15), maintaining the proper structure of tRNAs (16), modulating amino acid charging (17), and facilitating recognition by the ribosome (18). Despite their importance, our current understanding of chloroplast tRNA modifications is primarily limited to selected tRNAs in certain species (14). Traditionally, tRNA modifications were detected using liquid chromatography mass spectrometry (LC-MS). While LC-MS is sensitive and versatile, its application is restricted to selected tRNAs or provides a global view of modifications without precise information about the modification sites (19, 20). Consequently, sequencing-based methods, such as YAMAT-seq (21), QuantM-tRNAseq (22), mim-tRNAseq (23), or ALL-tRNAseq (24) (generally tRNA-seq), have been recently developed to offer genome-wide and nucleoside-specific insights into modified nucleosides. The detection of modified nucleosides using tRNA-seq is based on the ability of reverse transcriptase (RT) to introduce mismatches in synthesised cDNA at sites corresponding to modified nucleosides in the RNA or to block reverse transcription at modified nucleosides. By mapping mismatches in the cDNA or RT stops, one can deduce the locations of the RNA modifications. Importantly, the mismatches contain an additional layer of information, as the misincorporated nucleosides are not random and can reflect the chemical nature of the modified nucleoside; thus, misincorporation signatures can be used to distinguish modifications (23). tRNA poses a challenge for reverse transcription as a significantly high proportion of its nucleosides are modified, and its stable structure can inhibit RT. However, careful selection of proper RT and reverse transcription conditions has proven effective in analysing tRNAs and their modifications in diverse organisms (23, 24). Notably, not all types of chemical modifications can be detected using tRNA-seq, and this method mainly detects modifications occurring on the Watson-Crick face. Therefore, we decided to utilise both tRNA-seq and LC-MS to provide an overview of tRNA modifications in chloroplasts.

Additionally, we analyse publicly available Arabidopsis RNA sequencing data, focusing on detecting nucleoside modifications through chemical conversion and increased misincorporation frequency by reverse transcriptase at modified nucleosides. Furthermore, the homology between prokaryotic and organellar systems allows us to identify known and putative sites of chloroplast rRNA modifications. To achieve this, we utilise the secondary RNA structure and the cryo-EM structures of the chloroplast ribosome from *Spinacia oleracea*, along with the significant sequence similarity between *S. oleracea* and *Arabidopsis thaliana* rRNAs (3–6, 25, 26).

The present study aims to consolidate information on tRNA and rRNA modifications within chloroplasts. Our analysis confirms several modifications at positions corresponding to prokaryotic counterparts and identifies potential novel modified nucleosides in chloroplast tRNAs and rRNAs, possibly specific to plants. Furthermore, we discuss the molecular roles of these described modifications and their potential implications for cell biology.

## MATERIALS AND METHODS

### tRNA-seq library preparation

#### Total RNA isolation

Total RNA was isolated with 1 mL home-made Trizol substitute (27) from 100 mg ground tissue of 3.5-week Arabidopsis Col-0 leaves (grown in long-day photoperiod 16 h light/8h dark, 140 µmol*m^-2^*s^-1^, 23°C/21°C) or 2 mL log phase culture of *E. coli* str. K-12 substr. MG1665 (OD_600_ = 0.6). Phase separation was achieved with the addition of 0.2 mL chloroform. This was followed by two additional purifications of the aqueous phase using homemade Trizol and chloroform (0.65 mL and 0.13 mL for the first round, 0.325 mL and 0.07 mL for the second round). RNA was precipitated with 2.5 volumes of 99.6% ethanol overnight at −80°C. The resulting pellet was air-dried and resuspended in nuclease-free water.

#### tRNA isolation

The tRNA fraction was isolated from 17 µg of total RNA using Nucleobond AX-20 columns (Macherey-Nagel) following a rescaled protocol based on (28). The tRNA was then deacylated with 50 mM Tris-HCl (pH 9.0) for 45 minutes at 37°C, dephosphorylated using 10 U T4 PNK (ThermoFisher) for mim-tRNAseq libraries, and precipitated with 99.6% ethanol as per the mim-tRNAseq protocol (23). The concentration of tRNA was determined using the Qubit RNA HS Assay, and 165 ng was used for adapter ligation.

#### Adapter ligation

For mim-tRNAseq libraries, 3’ adapters were ligated according to (23). For QuantM-tRNAseq libraries, 5’ and 3’ adapters were ligated as described in (22), with an incubation period of 3 hours.

#### Reverse transcription

Reverse transcription was performed with Induro (New England Biolabs) following the manufacturer’s instructions. Adapter-tRNA, RT primer, and dNTPs were mixed, incubated at 65°C for 5 minutes, and then placed on ice. The remaining components were added, and the 20-µL reaction was incubated at 50°C for 2 hours. This was followed by the addition of 1 µL of 5 M NaOH and a 3-minute incubation at 95°C. For reverse transcription with SuperScript IV (ThermoFisher), adapter-tRNA was mixed with RT primer and incubated at 82°C for 2 minutes, 25°C for 5 minutes, and then placed on ice. The following components were then added: 1x SSIV-Mn^2+^ buffer (50 mM Tris-HCl pH 8.3, 50 mM KCl, 4 mM MnCl_2_), 5 mM freshly prepared DTT, 200 U SuperScript IV, and 8 U RNase Inhibitor Murine (New England Biolabs). The reaction was incubated at 50°C for 10 minutes, followed by the addition of 0.5 mM dNTPs, and incubation continued for 2 hours. The reaction was further incubated at 80°C for 10 minutes and at 95°C for 3 minutes after the addition of 1 µL of 5 M NaOH.

#### Circularization, amplification and sequencing

Gel-purified cDNAs were circularised using CircLigase ssDNA Ligase (Lucigen) and 5 M betaine (VWR) according to the mim-tRNAseq protocol (23). Libraries were amplified with Q5 Polymerase (New England Biolabs) with an initial denaturation at 98°C, followed by 11-13 cycles of 10 seconds at 98°C, 15 seconds at 65°C, and 20 seconds at 72°C. Libraries were resolved on a 6% PAGE gel, and 130-230 bp (mim-tRNAseq) or 140-260 bp (QuantM-tRNAseq) fragments were excised and eluted as per the mim-tRNAseq protocol ((23)). The concentration was estimated using the NEBNext Library Quant Kit for Illumina (New England Biolabs). Each 10 nM library pool (mim-tRNAseq or QuantM-tRNAseq) was sequenced for 2×100 cycles on the Illumina NovaSeq 6000 platform in separate flow cells (650M clusters in total).

### tRNA-seq read processing and data analysis

In the first step, the R1 and R2 reads were merged using BBMap tool with the following parameters: `bbmerge.sh adapter1=AGATCGGAAGAGCACACGTCTGAACTCCAGTCAC adapter2=AGATCGGAAGAGCGTCGTGTAGGGAAAGAGTGT mininsert=25 mininsert0=25 vstrict=t`. Then, adapters were removed using cutadapt v4.9. For QuantM-tRNAseq, the parameters were: `cutadapt -a GTATCCAGTX -u 2 and cutadapt -g XCAGAGTTCTACAGTCCGACGATC -q 25`. For mim-tRNAseq parameters were: `-u −10 -u 2`. Next, the mim-tRNAseq v1.3.8 pipeline was used to analyse *A. thaliana* (mapping with our genome) and *E. coli* reads (mapping with built-in genome) using the following parameters: `mimseq --cluster-id 0.99 --threads 15 --min- cov 0.0005 --max-mismatches 0.1 --max-multi 4 --remap --remap-mismatches 0.075`. The mimseq output files containing misincorporations and hard stop positions, were used for further analysed and visualised using R v4.3.3 R Core Team, 2024).

### RT misincorporation assay for rRNA modifications (cRT-PCR and amplicon sequencing)

The positions of chloroplast rRNA modifications were determined in various RT misincorporation assays. Firstly, the modification in a fragment of 16S rRNA (m^2^G915) was detected using TGIRT-III (InGex) in recently published datasets (29, 30). For a broader analysis of 16S and 23S rRNA modifications, we employed two approaches – Sanger sequencing of amplicons and plasmids with cloned cRT-PCR products.

Total RNA was isolated using a ‘homemade’ Trizol substitute as described in (27) from 100 mg of ground tissue from 3.5-week-old Arabidopsis Col-0 leaves. Following DNase I treatment, 2 µg of total RNA was circularised in a reaction containing 6 U T4 RNA Ligase 1 (New England Biolabs), 50 µM ATP, 1x T4 RNA ligase buffer, and 4 U RNase Inhibitor Murine, incubated at 37°C for 1 hour and then at 65°C for 15 minutes. The circularised RNA (cRNA) was precipitated with 99.6% ethanol. One microgram of cRNA was used for reverse transcription with 2 pmoles of a 16S rRNA-specific primer (Supplementary Table 8) and SuperScript II (ThermoFisher), following the manufacturer’s instructions. The cDNA was amplified using Q5 Polymerase (New England Biolabs) with an initial denaturation at 98°C for 30 seconds, followed by 30 cycles of 98°C for 30 seconds, 67°C for 20 seconds, and 72°C for 18 seconds, with a final extension at 72°C for 2 minutes. The resulting amplicons were cloned into the pMiniT2.0 vector (New England Biolabs), and plasmid DNA from 18 colonies was purified and Sanger sequenced. For amplicon Sanger sequencing, DNase I-treated total RNA from Arabidopsis was used for reverse transcription with Induro (New England Biolabs) and random hexamers, following the manufacturer’s instructions. The resulting cDNA and qDNA were amplified with Q5 Polymerase (New England Biolabs), and the gel-purified PCR products were sequenced.

### m^5^C analysis

We utilised publicly available Arabidopsis data from (31) to detect m^5^C in chloroplast rRNAs. The SRA files (SRR3347461, SRR3347462, SRR3347463, and SRR3347464) were downloaded from the NCBI Sequence Read Archive using the prefetch function from SRAtoolkit (version 3.0.6). Next, the files were converted to .fastq format using fasterq-dump and compressed with gzip using pigz (version 2.4). Subsequently, adapter sequences were removed, and quality trimming was performed using trim_galore! (version 0.6.10) with the following arguments: `-j 8 -q 30 --three_prime_clip_R1 1 -- three_prime_clip_R2 1 --clip_R1 11 --clip_R2 11 --paired`. The paired-end, trimmed reads were then mapped to the *A. thaliana* rRNA genes with Cs converted to Ts using hisat-3n (2.2.1-3n-0.0.3) with the following arguments: `--base-change C,T -- no-spliced-alignment --bowtie2-dp 1 --no-unal --directional-mapping-reverse -p 16`. The bisulfite unconverted Cs (i.e., m^5^C or m^4^C) were identified next using the pileup() function from the Rsamtools package (version 2.16.0). Positions with a minimum of 200 mapped reads were analysed.

### m^7^G analysis

The m^7^G presence in chloroplast rRNAs and tRNAs was analysed based on publicly available data published by (32). SRA files representing long reads (e.g. rRNAs, SRR8130784 SRR8130785) and short reads (e.g. tRNAs, SRR8130786 SRR8130787 SRR8130788 SRR8130789 SRR8130790 SRR8130791) were downloaded from the NCBI Sequence Read Archive and converted to .fastq format as described above. The analysis was conducted separately for long and short reads, following the methodology outlined in (32), utilising the scripts provided by the authors of the original publication (https://github.com/jeppevinther/m7g_map_seq/blob/master/m7g_map_seq.sh) with minor modifications as described below. The long RNAs were sequenced in paired-end mode; thus, adaptor sequences and quality trimming were performed for each read from the pair using cutadapt (version 2.8) with the following options: `-a AGATCGGAAGAGCACACGTCT -A AGATCGGAAGAGC --nextseq-trim=20 -j 12`. Subsequently, barcodes were removed from the reads using the preprocessing.sh script from the RNAProbR package (33). In the next step, reads without barcodes were mapped to the *A. thaliana* rRNA genes using bowtie2 (version 2.3.5.1) with the following arguments: `-p 12 --local -N 1 -D 20 -R 3 - L 15`. PCR duplicates were then removed based on the mapping location and the presence of identical barcodes using the collapse.sh script (https://people.binf.ku.dk/jvinther/data/RNA-seq/collapse.sh). To detect m^7^G, the mpileup function from samtools (version 1.10), along with the getFreq2000.R script (https://github.com/jeppevinther/m7g_map_seq), was employed. Analysis of the short reads (sequencing performed in single-end mode) followed a similar procedure but without barcode removal and collapsing of duplicated reads. For rRNAs analysis, sites with a minimum coverage of 500 reads were analysed, while for tRNAs, sites with a minimum coverage of 1500 reads were considered.

### Analysis of rRNA modifications using RT

#### UPLC-MS analysis of tRNA modifications

Following total RNA extraction, tRNA purification was performed as previously described (34) with minor changes. Briefly, the total RNA was diluted in 10 ml of equilibration buffer EQ (10 mM Tris-HCl, pH 6.3, 15% ethanol, 200 mM KCl) prior to purification on a Nucleobond AX-100 column (Macherey-Nagel). The column-bound RNA was washed twice with wash buffer WB (10 mM Tris-HCl, pH 6.3, 15% ethanol, 400 mM KCl) and the tRNA containing fraction was eluted with elution buffer EB (10 mM Tris-HCl, pH 6.3, 15% ethanol, 750 mM KCl) into 2.5 vol. of 99.6% ethanol. The tRNA was precipitated over two nights at −20 °C and pelleted by centrifugation at 10 000 *g*, 4 °C for 30 min. The pellet was washed with 80% ethanol to remove excess salt. The final tRNA pellet was air-dried at room temperature and resuspended in RNase/DNase-free water. For UPLC-MS analysis, the purified tRNA was mixed with m^1,3^Ψ (internal standard; 25 ng of m^1,3^Ψ per 1 µg of tRNA) and cleaved and dephosphorylated into mononucleosides as previously described (35). Modified ribonucleosides were separated on an Acquity UPLC® BEH (Ethylene Bridged Hybrid) C18 column (Ø 1.7 μm, 2.1 × 150 mm, Waters) coupled to an ExionLC UPLC with AB Sciex QTRAP-6500+ detector. The ribonucleosides were analysed by multiple reaction monitoring (MRM) using ESI+ mode according to the previously developed method (34). Please refer to Supplementary Table 7 for MRM parameters for the modified ribonucleosides.

## RESULTS AND DISCUSSION

### Utilising tRNA-seq to detect modifications

Given the shared prokaryotic origin and similar expression machinery of chloroplasts and prokaryotes (2), we included *E. coli*, a well-established model for tRNA modifications (36), alongside Arabidopsis leaf samples in our experiment. We took a systematic approach, testing two RTs: Induro and SuperScript IV. We optimised reverse transcription conditions and compared two tRNA-seq pipelines (mim-tRNAseq and QuantM-tRNAseq). Each sample was analysed in triplicate (details in Experimental Procedures). Our method involved isolating tRNAs, followed by adapter ligation, reverse transcription, library amplification, sequencing, and bioinformatic analysis (Figure 1a).

**Figure 1.**
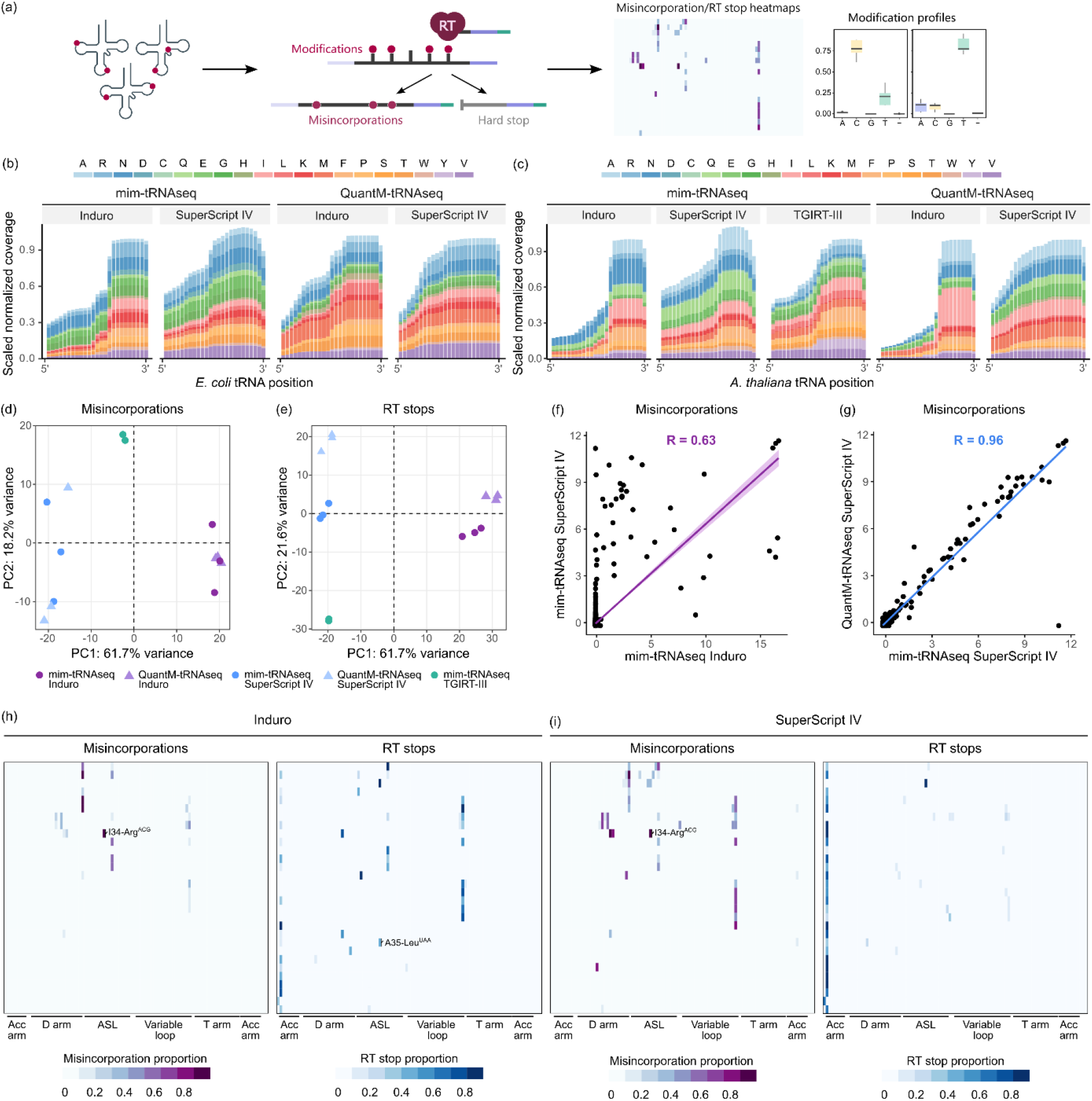
tRNA-seq reveals multiple modified nucleosides in chloroplast tRNAs. (**a**) Schematic representation of the tRNA-seq workflow and identification of modified nucleosides based on reverse transcriptase (RT) misincorporations and stalling events (RT stops). (**b**, **c**) Scaled coverage of sequencing reads for tRNA isodecoders from *E. coli* (**b**) and Arabidopsis chloroplasts (**c**). Data are presented for two tRNA-seq library preparation methods (mim-tRNAseq and QuantM-tRNAseq) and two reverse transcriptases (SuperScript IV and Induro). For comparison, sequencing data from (Arrivé et al., 2023) using TGIRT-III reverse transcriptase is included. (**d**, **e**) Principal component analysis (PCA) of RT misincorporations (**d**) and RT stops (**e**). Each point represents an individual biological replicate, grouped by colour and shape according to the tRNA-seq pipeline and RT used. (**f**, **g**) RT misincorporation correlation plots: (**f**) compares misincorporations between Induro and SuperScript IV RTs, while (**g**) shows the correlation between mim-tRNAseq and QuantM-tRNAseq pipelines using SuperScript IV. Pearson’s correlation coefficient (R) values are shown for each plot. (**h, i**) Heatmaps illustrating the frequency of RT misincorporations (top) and RT stops (bottom) for Induro (**h**) and SuperScript IV (**i**) RTs across chloroplast tRNAs. Abbreviations: Acc arm – Acceptor arm, ASL – Anticodon stem-loop.

We generated 24 sequencing libraries and sequenced them on an Illumina platform using 2×100 bp paired-end reads, allowing for complete tRNA coverage. Each library yielded 17-31 million reads, with 2.5-23 million mapping to tRNA sequences (Supplementary Table 5). Arabidopsis samples specifically showed 0.23-2.5 million reads mapping to chloroplast tRNAs (8-22%; Supplementary Table 6). The fraction of reads mapped to cytosolic tRNAs ranged from 78 to 92%, while reads mapped to mitochondrial tRNAs ranged between 0.12-0.3%. These data are consistent with the high translational activity of chloroplasts in leaves (37).

tRNA coverage after alignment varied significantly between samples (Figure 1b, c). Induro RT generated a bias towards the 3’ end, while SuperScript IV provided more even coverage. *E. coli* tRNAs showed better coverage (Figure 1b) compared to Arabidopsis chloroplast tRNAs (Figure 1c), suggesting potentially more RT-blocking modifications in chloroplast tRNAs. Interestingly, our data (generated using SuperScript IV) mirrored coverage patterns observed in a recent study using TGIRT-III and mim-tRNAseq (Figure 1c), supporting the notion that non-ideal coverage can be caused by chloroplast tRNA properties. Notably, Induro RT introduced frequent RT stops, leading to higher coverage at the 3’ end that gradually decreased towards the 5’ end. We identified SuperScript IV with QuantM-tRNAseq as the combination providing the most even coverage for chloroplast tRNAs (Figure 1c). Similarly, this combination of RT and tRNA-seq pipeline provided the most even coverage of *E. coli* tRNAs (Figure 1b). Importantly, we observed a balanced representation of each chloroplast tRNA isodecoder, allowing for faithful analysis of every tRNA.

Next, we compared similarities between samples in terms of RT misincorporations and RT stops (Figure 1d-g). We observed a high similarity of RT misincorporations between biological replicates (Figure 1d), with even higher similarity of RT stops (Figure 1e). Notably, the main factor that differentiated samples was the type of RT used, with the impact of the tRNA-seq pipeline being significantly lower. Consistently, the correlation of RT misincorporations between individual samples generated by Induro RT and SuperScript IV was significantly lower (R = 0.63, Figure 1f) than between two samples generated by two tRNA-seq pipelines with the same RT (R = 0.96, Figure 1g). In summary, we observed high similarity between biological replicates in both RT misincorporations and RT stops. Importantly, the sequencing depth providing sufficient coverage of tRNAs and high similarity within replicates allowed us to analyse tRNA modifications for all Arabidopsis chloroplast tRNAs and compare them to *E. coli*.

Next, we generated RT misincorporation and RT stop profiles across all 30 chloroplast tRNAs (Figure 1h, i) and *E. coli* tRNAs (Supplementary Figure S2). Consistent with tRNA coverage analysis (Figure 1b, c), more RT stops were generated with Induro RT than SuperScript IV. However, some RT stops, such as at C34 in tRNA-Ile^CAU^ (discussed below), were consistently introduced by both RTs. Conversely, SuperScript IV generated significantly more RT misincorporations than Induro RT (Figure 1h, i, Supplementary Figure S2). Both types of RT marks reflect the sites of modified nucleosides, and we observed that most of them can be found in the anticodon arm, at position 26, and at position 47, consistent with commonly found sites of modifications in prokaryotic tRNAs (38, 39). Although little is known about the tRNA modifications in Arabidopsis chloroplasts, we could detect the described inosine at the wobble position in tRNA-Arg^ACG^ (39, 40) in RT misincorporation profiles (Figure 1h, i). Notably, the profiles of RT marks differed significantly among individual tRNAs. For example, we observed an increased Induro RT’s hard stop signal at chloroplast position 35 only in the tRNA-Leu^UAA^ (Figure 1h, Supplementary Figure S3a), suggesting a modification at wobble U34. This observation is in accordance with the fact that *E. coli* tRNA-Leu^UAA^ has cmnm^5^Um at position 34 (Supplementary Table 2) allowing for accurate decoding in the UUN split codon box. In some tRNAs, up to three modified nucleosides were detected (e.g., tRNA-Phe^GGA^), while there were also tRNAs without any detected modifications. Overall, chloroplast tRNAs had on average 1.2 detected modifications per tRNA compared to 1.3 in *E. coli* (Supplementary Tables 1 and 2). To sum up, we detected many modified nucleosides in chloroplast tRNAs that were not described earlier.

### Identifying tRNA modifications at position 37 in chloroplasts

Nucleoside 37, located directly downstream of the anticodon in chloroplast tRNAs, is always a purine base (adenine or guanine). This position is frequently modified in various systems, especially when the first codon position is A or U. These modifications stabilise codon-anticodon interactions through base stacking and prevent interaction with U33, keeping the anticodon stem-loop open for codon recognition. Consequently, the absence of a modification at position 37 can increase the risk of frameshifting errors during translation.

In prokaryotes, common modifications at this position include 2-methylthio-*N*^6^-isopentenyladenosine (ms^2^i^6^A), *N*^6^-threonylcarbamoyladenosine (t^6^A), *N*^6^-methylthreonylcarbamoyladenosine (m^6^t^6^A), *N*^6^-methyladenosine (m^6^A), 2-methyladenosine (m^2^A), and 1-methylguanosine (m^1^G). Recently, the presence of m^2^A37 has been documented in four chloroplast tRNAs: tRNA-Arg^ACG^, tRNA-His^GUG^, tRNA-Met^eCAU^, and tRNA-Ser^GGA^ (Supplementary Table 1, (41)). This modification facilitates efficient translation by maintaining the tRNA anticodon stem-loop in an open conformation (41). However, the potential for other modifications at this position in chloroplast tRNAs remains unclear.

We began by examining the types of modifications detectable by Induro and SuperScript IV RTs in the well-characterised system, *E. coli*. Similar to chloroplasts, *E. coli* also has exclusively purines at position 37 (Supplementary Table 2). We observed that six A37 positions induced RT misincorporations and RT stops in *E. coli* (Figure 2a, b). Notably, Induro was more susceptible to blockage, while SuperScript IV introduced misincorporations at A37 (Figure 2b). Importantly, all A37 positions leaving RT marks are known to possess ms^2^i^6^A, suggesting that both RTs can detect this modification. While both enzymes generated similar ms^2^i^6^A signatures, Induro produced a more consistent pattern across analysed tRNAs (Figure 2c). Importantly, other adenosine modifications at position 37 in *E. coli* (m^2^A, t^6^A, i^6^A, m^6^t^6^A, and m^6^A) did not generate RT stops or misincorporations in our datasets, suggesting that they could not be detected by tRNA-seq in this context. *E. coli* also possesses eight tRNAs with G37, all modified to m^1^G. Notably, all these m^1^G37 tRNAs generated RT marks (Figure 2d, e), along with specific misincorporation signatures depending on the RT used (Figure 2f). This confirms that m^1^G can be detected by both enzymes. In conclusion, we have demonstrated that tRNA-seq using Induro and SuperScript IV RTs can readily detect two common modifications at position 37: ms^2^i^6^A and m^1^G.

**Figure 2.**
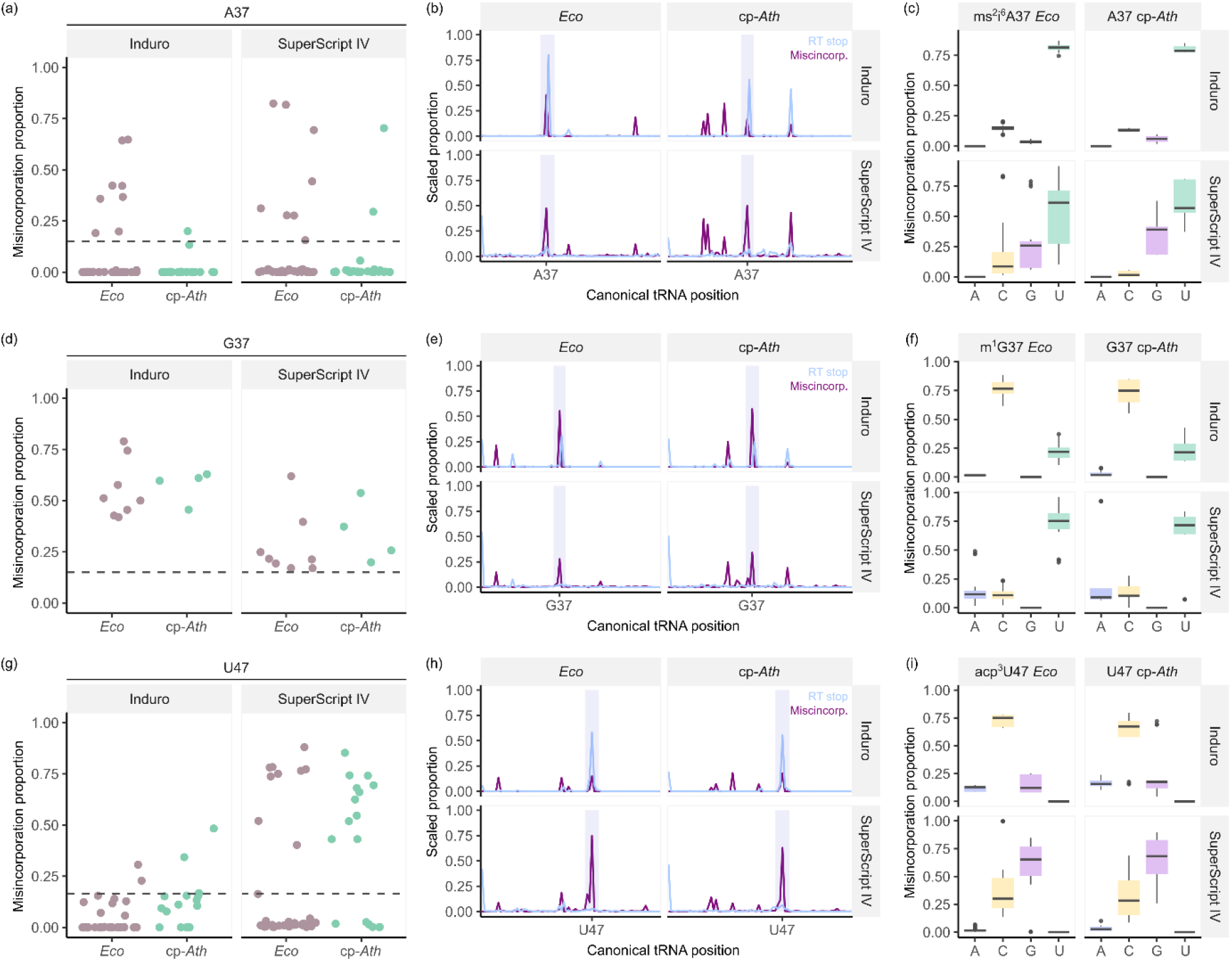
tRNA-seq reveals similarities in tRNA modifications between *E. coli* and Arabidopsis chloroplasts. (**a**, **d**, **g**) Dot plots depicting the proportion of RT misincorporations for specific nucleosides: (**a**) A37, (**d**) G37, and (**g**) U47. Data are presented for *E. coli* (light brown) and Arabidopsis chloroplasts (green), using both Induro (left) and SuperScript IV (right) RTs. Points above the dashed line (RT misincorporation proportion > 0.15) likely represent modified nucleosides in these tRNAs. (**b**, **e**, **h**) Line plots showing RT misincorporations (purple) and RT stops (blue) for *E. coli* (left) and Arabidopsis chloroplasts (right) generated with both Induro (top) and SuperScript IV (bottom) RTs. These plots depict average values across all tRNAs exceeding the threshold (as in panels **a**, **d** and **g**) for a specific RT or organism. The analysed sites with RT misincorporation proportions exceeding the given threshold are highlighted in light grey. (**c**, **f**, **i**) RT misincorporation signatures for the same set of tRNAs analysed in panels (**b**, **e**, **h**). These signatures show the frequency of misincorporated nucleosides by Induro (top) or SuperScript IV (bottom) RTs at specific nucleoside positions in tRNAs from *E. coli* (left) and Arabidopsis chloroplasts (right).

We next investigated whether ms^2^i^6^A and m^1^G could be detected in chloroplast tRNAs. Three chloroplast tRNAs with A at position 37 (tRNA-Phe^GAA^, tRNA-Ser^UGA^, and tRNA-Tyr^GUA^) generated RT misincorporations and RT stops (Figure 2a,b). Importantly, the misincorporation signatures closely resembled those generated at ms^2^i^6^A in *E. coli*, suggesting the presence of ms^2^i^6^A in these chloroplast tRNAs (Figure 2c). To further confirm the presence of ms^2^i^6^A, we used liquid chromatography-mass spectrometry (LC-MS) on total tRNA extracts. The results identified ms^2^i^6^A, supporting its existence in chloroplast tRNAs (Supplementary Table 7). Since ms^2^i^6^A is specific to prokaryotic systems, it is assumed that in plant cells it can be found in chloroplast and mitochondrial tRNAs. Given the significantly higher abundance of chloroplast tRNAs compared to mitochondrial tRNAs in leaf tissue (Supplementary Table 6), it is likely that the observed ms^2^i^6^A is predominantly present in chloroplast tRNAs. Next, we analysed chloroplast tRNAs with G at position 37 (tRNA-Gln^UUG^, tRNA-Leu^UAG^, tRNA-Pro^UGG^, and tRNA-Val^GAC^). Interestingly, all these tRNAs displayed RT stops and misincorporations (Figure 2d, e) with signatures highly similar to the m^1^G37 signatures observed in *E. coli* (Figure 2f). These results strongly suggest the presence of both ms^2^i^6^A and m^1^G modifications at position 37 in chloroplast tRNAs.

### Modifications within the chloroplast variable loop exhibit similarities to those found in *E. coli*

*E. coli* tRNAs contain two types of modified nucleosides within the variable loop: 7-methylguanosine (m^7^G) at position 46 and 3-(3-amino-3-carboxypropyl)uridine (acp^3^U) at position 47 (Supplementary Table 2). Both modifications significantly impact tRNA stability. Specifically, m^7^G forms a tertiary base pair with C13 and U22, introducing a positive charge that stabilises the tRNA’s tertiary structure (42). Interestingly, m^7^G also promotes the formation of acp^3^U at position 47 (43). acp^3^U47, a highly conserved modification in prokaryotes, enhances the thermal stability of tRNA by increasing its overall structural stability (44).

Enroth *et al.* (32) provided insights into detecting m^7^G in rRNAs and tRNAs, including those in Arabidopsis chloroplasts. The authors used m^7^G mutational profiling sequencing (m^7^G-MaP-seq) to identify m^7^G modifications at the nucleoside level. This method employed a sodium borohydride (NaBH_4_) reagent to convert m^7^G-modified nucleosides into abasic sites, subsequently detected as misincorporations in cDNA through reverse transcription and sequencing. Their data allowed us to map m^7^G in chloroplast tRNAs, identifying seven chloroplast tRNAs with this modification at position 46 (Supplementary Figure 2b).

In *E. coli*, acp^3^U47 is found in multiple tRNAs (Supplementary Table 2). As acp^3^U affects RT (23), we identified RT marks at U47 for ten tRNAs (Figure 2g,h). The acp^3^U47 generated RT-specific misincorporation signatures (Figure 2i). Similarly, U47 in chloroplast tRNAs is commonly modified (Figure 2g,h). The RT misincorporation signatures in chloroplast tRNAs highly resembled those of acp^3^U47 in *E. coli* (Figure 2i), suggesting that uridines at position 47 in chloroplast tRNAs are also modified to acp^3^U47. Collectively, these data indicate that modifications within the chloroplast variable loop are similar to those found in *E. coli*, which are crucial for stabilising the tRNA’s tertiary structure.

### RT misincorporation signatures reveal a putative novel modification in Arabidopsis chloroplast tRNA^Ile^(CAU)

Chloroplasts utilise two distinct tRNA molecules to decode isoleucine codons (AUH, where H represents A, C, or U). The prevalent AUU and AUC codons are recognised by tRNA-Ile^GAU^. However, decoding the rare AUA codon presents a unique challenge. This specific codon is exclusively recognised by tRNA-Ile^CAU^, which harbours a crucial modification at its wobble position (C34) – a cytidine residue. This modification ensures fidelity by preventing misreading of AUG codons, which encode methionine, by tRNA-Ile^CAU^ (43). In bacteria and archaea, the wobble cytidine undergoes similar modifications, either to 2-lysylcytidine (lysidine, k^2^C) or 2-agmatinylcytidine (agmatidine), respectively. Interestingly, distinct enzyme classes catalyse these modifications despite their chemical similarities (45). While current research suggests lysidine as the likely modification in chloroplast tRNA-Ile^CAU^ (46, 47), experimental validation is necessary.

We investigated tRNA-Ile^CAU^ from *E. coli*, a well-characterised system with known lysidine modification, and compared it to the equivalent tRNA in Arabidopsis chloroplasts. *E. coli* encodes two tRNA-Ile^CAU^ isodecoders, with at least one possessing lysidine at C34 (48). Lysidine, a cytidine derivative with a lengthy lysine moiety, potentially blocks reverse transcription, enabling its detection. As expected, lysidine blocked both employed RTs in *E. coli* (Figure 3a, b). Interestingly, a significant blockage of these RTs also occurred at C34 in chloroplast tRNA-Ile^CAU^, supporting the notion of a modification at this position. Notably, Induro RT was entirely blocked by lysidine in *E. coli* and the C34 modification in Arabidopsis, while SuperScript IV displayed partial blockage but continued cDNA synthesis with visible misincorporations (Figure 3a, b). Therefore, we compared the misincorporation signatures generated by SuperScript IV at C34 of tRNA-Ile^CAU^ in *E. coli* and Arabidopsis chloroplasts. In *E. coli*, k^2^C34 resulted in similar incorporations of A, G, and T in place of the modified cytidine (Figure 3c). Surprisingly, C34 in Arabidopsis tRNA-Ile^CAU^ produced a distinct signature. Here, C34 was primarily read as A with a minor proportion of U, completely lacking G.

**Figure 3.**
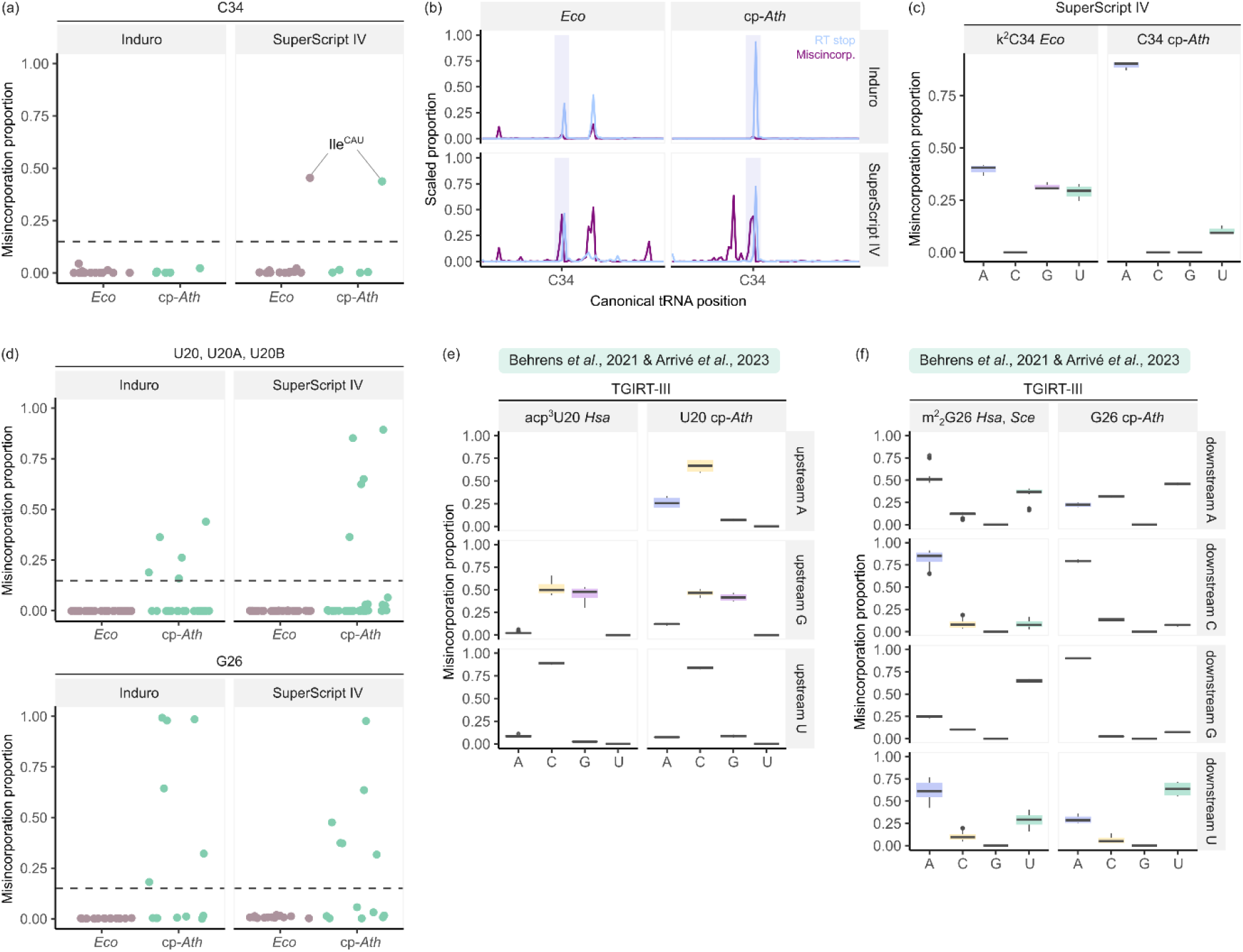
Chloroplast tRNAs exhibit specific modifications distinct from *E. coli*. (**a**) Dot plots depicting the proportion of RT misincorporations at cytidine C34 in *E. coli* (light brown) and Arabidopsis chloroplasts (green). Both organisms exhibit a modified cytidine at this position, as indicated by points exceeding the threshold (dashed line, RT misincorporation proportion > 0.2). Notably, only SuperScript IV RT introduces misincorporations at C34. (**b**) Line plots showing RT misincorporations (purple) and RT stops (blue) for *E. coli* (left) and Arabidopsis chloroplasts (right) generated with both Induro (top) and SuperScript IV (bottom) RTs. Analysed sites exceeding the threshold from panel (**a**) are highlighted in light grey for tRNA-Ile^CAU^ in both *E. coli* and Arabidopsis chloroplasts. (**c**) RT misincorporation signatures for tRNA-Ile^CAU^, known to possess lysidine (k^2^C34) at position 34 in *E. coli* (43). These signatures depict the frequency of misincorporated nucleosides by SuperScript IV RT at C34 in tRNA-Ile^CAU^ from *E. coli* (left) and Arabidopsis chloroplasts (right). Note the distinct misincorporation signatures between the two organisms, suggesting potential differences in the modification at this site. (**d**) Dot plots depict the proportion of RT misincorporations for U8 (left) and G26 (right) using the same colour scheme and layout as panel (**a**). (e) RT (TGIRT-III) misincorporation signatures for annotated m^2^_2_G26 in *Saccharomyces cerevisiae* and *Homo sapiens* tRNAs (data from (23); right) and two groups of *Arabidopsis thaliana* G26 (data from (52); left). (**f**) Sodium borohydride treatment and mutational profiling were used to detect *N*^7^-methylguanosine (m^7^G) (Enroth *et al.*, 2019). This panel presents a plot of mutation rate difference (x-axis) versus −10*log(p-val) (y-axis) for all Arabidopsis chloroplast tRNAs.

Previous studies documented that misincorporation signatures can vary significantly depending on the surrounding sequence context for certain modifications (23). This variation could explain the observed differences between *E. coli* and Arabidopsis. However, our analysis revealed that the sequence context surrounding C34 is identical in the analysed tRNAs (Supplementary Figure S4). Therefore, the contrasting misincorporation signatures between *E. coli* and Arabidopsis chloroplasts at C34 in tRNA-Ile^CAU^ suggest the presence of a modification at the wobble C in Arabidopsis tRNA-Ile^CAU^ that differs from lysidine.

To our knowledge, the only prior indication of a potential modification at C34 in chloroplast tRNA-Ile^CAU^ comes from a study by Francis and Dudock (49) analysing spinach, although the specific modification was not identified. Interestingly, the Arabidopsis orthologue of the *E. coli* enzyme (TilS) responsible for lysidine modification remains undiscovered. Similarly, the orthologue of TiaS, the enzyme catalysing agmatidine formation in archaeal tRNA-Ile^CAU^, has not been found in Arabidopsis. In conclusion, our findings strongly suggest a modification at C34 in tRNA-Ile^CAU^ isolated from Arabidopsis chloroplasts. However, the exact nature of this modification remains elusive.

### Non-prokaryotic modification subset around D-arm of chloroplast tRNAs

The region of the D-arm and its adjacent nucleosides is marked by modifications enhancing the cloverleaf and L-shaped structure of tRNA (50). It also determines recognition by aminoacyl-tRNA synthetases (51). The prevailing dihydrouridine appears in the D-loop, while the other modification types mainly occur in the stem and the neighbouring junctions with other tRNA parts. Our sequencing data revealed several distinctions between the modification subset of prokaryotic and chloroplast tRNAs. *E. coli* tRNAs contain 4-thiouridine (s^4^U) at positions 8 and 9, 2’O-methylguanine (Gm) at position 18, and dihydrouridine (D) at positions 16, 17, 20, 20a and 21. Since only s^4^U out of the mentioned modifications occurs in the Watson-Crick face, it could be detected by our method. Both RTs induced elevated levels of misincorporation at U8, which was distinctive for *E. coli* tRNAs and did not appear in chloroplast tRNAs (Supplementary Figure S3c).

Simultaneously, we observed an increased misincorporation rate at U20,20a,20b and G26 in chloroplast tRNAs exclusively (Figure 3d). Given the presence of other modification types in the positions, we decided to extend our analysis of the misincorporation signature comparison to eukaryotic tRNAs. Thus, we employed mim-tRNAseq data from *Homo sapiens* and *Saccharomyces cerevisiae* (23) and Arabidopsis (52), both generated with TGIRT-III reverse transcriptase (Figure 3e, f).

In humans, several tRNAs are modified at U20 and U20a into acp^3^U (23, 44). The only known function of acp^3^U20a is the coordination of Mg^2+^, promoting the local tRNA conformation and efficient translation (53). As demonstrated earlier, acp^3^U induces RT misincorporations and appears at U47 of chloroplast tRNAs (Figure 2g-i). Behrens *et al.* (23) showed that the acp^3^U modification mark of TGIRT-III significantly depends on the upstream sequence context. Notably, the chloroplast U20,20a,20b signatures resemble the human acp^3^U20,20a marks within the available upstream contexts (Figure 3e). This data suggests the existence of eukaryote-specific tRNA acp^3^U modification in the D-arm of chloroplast tRNAs.

Modified G26 is highly conserved in eukaryotes, archaea, mitochondria and hyper-thermophilic bacteria (54–56). The residue is mono- or dimethylated at *N*^2^ of guanosine generating *N*^2^-methylguanosine (m^2^G) or *N*^2^,*N*^2^-dimethylguanosine (m^2^_2_G). The modification at G26 promotes binding with A/U44 in the variable arm, maintaining the tRNA tertiary structure (57). Although both modification types appear in nature, m^2^ G seems more efficient in pairing via 2 (instead of 3) hydrogen bonds in contrast to m^2^G, which leaves one hydrogen available. The feature is also reflected in RT’s modification reading manner. The detection of m^2^G using RT is unreliable, as only one out of 12 annotated sites at position 26 in human tRNAs could be identified. Consequently, we could determine the modification signature only for m^2^_2_G, which is independent of the sequence context in most cases (except on downstream G) (Figure 3f). The presented m^2^_2_G26 mark bases on previously annotated in MODOMICS and newly detected modifications in Behrens *et al.* (23). The G26 modification mark found in chloroplast tRNAs groups depending on the downstream sequence context and resembles m^2^ G26 marks for tRNAs with downstream C (Figure 3f). However, the signatures of chloroplast tRNAs with downstream A, G and U differ from those generated at m^2^ G26 in eukaryotes (Figure 3f). The modification marks are also not similar to the TGIRT-III’s m^1^G signature (23). Hence, these findings enforce the presence of m^2^_2_G26 only in a subset of chloroplast tRNAs. Considering that we do not know all the factors influencing modification reading way, we cannot exclude the presence of m^2^_2_G26 in all identified chloroplast tRNAs, decisively. Alternatively, the position could be modified differently, but it seems unlikely due to the high conservation of *N*^2^(,*N*^2^-di)methylguanosine across life domains. In conclusion, the data revealed the D-arm of chloroplast tRNAs is decorated with modifications which do not appear in this region in standard prokaryotic systems, such as *E. coli* (discussed below).

### The context of chloroplast tRNA modifications

The evolutionary established system of tRNA functionality maintenance highly depends on nucleoside modifications. Despite many tRNA modification functions being similar across life domains, a subset often varies significantly between species. Possible explanations for these differences include variations in molecular architecture and the characteristics of an organism’s habitat. For example, AU-rich mitochondrial tRNAs require m^1^A9 to avoid alternative, noncanonical structures, while m^5^s^2^U54 in *Thermus thermophilus* tRNAs is essential for protein synthesis at temperatures between 50-83°C (58, 59).

Given the similarity between prokaryotic and chloroplast molecular systems, one might expect the tRNA modification subset of *E. coli* to be reflected in chloroplasts. However, our results indicate only a partial resemblance, suggesting that the chloroplast translation machinery also employs derivative solutions found in eukaryotes and archaea. In this discussion, we consider the broader context of the (un)detected chloroplast tRNA modifications.

Modifications in the acceptor stem and D-arm interconnection include s^4^U8 and s^4^U9 (bacteria, archaea), m^1^G9, m^2^G10, m^2^_2_G10 (archaea, yeast, human), and m^1^A9 (human mitochondria). Prokaryotic s^4^U8 functions as a well-conserved tRNA structure stabiliser and near-UV light sensor. This residue pairs with A14 and increases the melting temperature of tRNA-Ser^GGA^ by 5°C (60). After UV treatment, it forms a cross-link bond with C13, leading to disruption of tRNA tertiary structure and translation inhibition (61, 62). Although s^4^U8 was detected in sequencing data of *E. coli* tRNAs, it was not found in chloroplasts (Supplementary Figure S3c, Figure 1h, i). Additionally, most of the mentioned modification types (except possibly m^2^G) in that region would induce RT misincorporations, if present in chloroplast tRNAs (23). Hence, only the presence of m^2^G remains a possibility. According to the MODOMICS database, *S. oleracea* chloroplast tRNA-Pro^UGG^ has a modified residue at G10 (14). It was suggested that the hydrophobic character of m^2^G10 enhances the structure of the D-loop (63). Taken together, we cannot definitively conclude that chloroplast tRNAs lack modifications at positions 8, 9 and 10. Nevertheless, the data suggests that modifications specific for prokaryotes, archaea and mitochondria (i.e. s^4^U, m^1^A, m^2^ G) are absent.

The other D-arm junction, with the anticodon arm, is noted for harbouring modifications at G26 and G27 in eukaryotes, hyperthermophilic bacteria and archaea. Modified G26 interacts with residues at positions 44 and 45, stabilising the hinge region, and occurs under specific sequence arrangements (Figure 4a, b; (64)). Interestingly, these requirements are diverse even within the same life domain. For example, yeast Trm1, which introduces m^2^G and m^2^_2_G, modifies tRNAs containing G10, C11 and at least five nucleosides in the variable loop. Conversely, yeast tRNA-Asp^GUC^, which does not meet any of these requirements, is monomethylated in wheat embryo extract and animal tissues (65–68). Chloroplast tRNA-seq revealed seven modified tRNAs at G26 into m^2^_2_G and/or m^2^G. Considering the nucleosides at positions 10 and 25, and the variable loop length, it is possible to observe similar dependencies in detected chloroplast tRNAs (G10:C25 or C10:G25) as found in the other species (66, 69–72). Therefore, the tRNA recognition factors for chloroplast Trm1 ortholog and their evolutionary context appear to be an interesting direction to explore.

**Figure 4.**
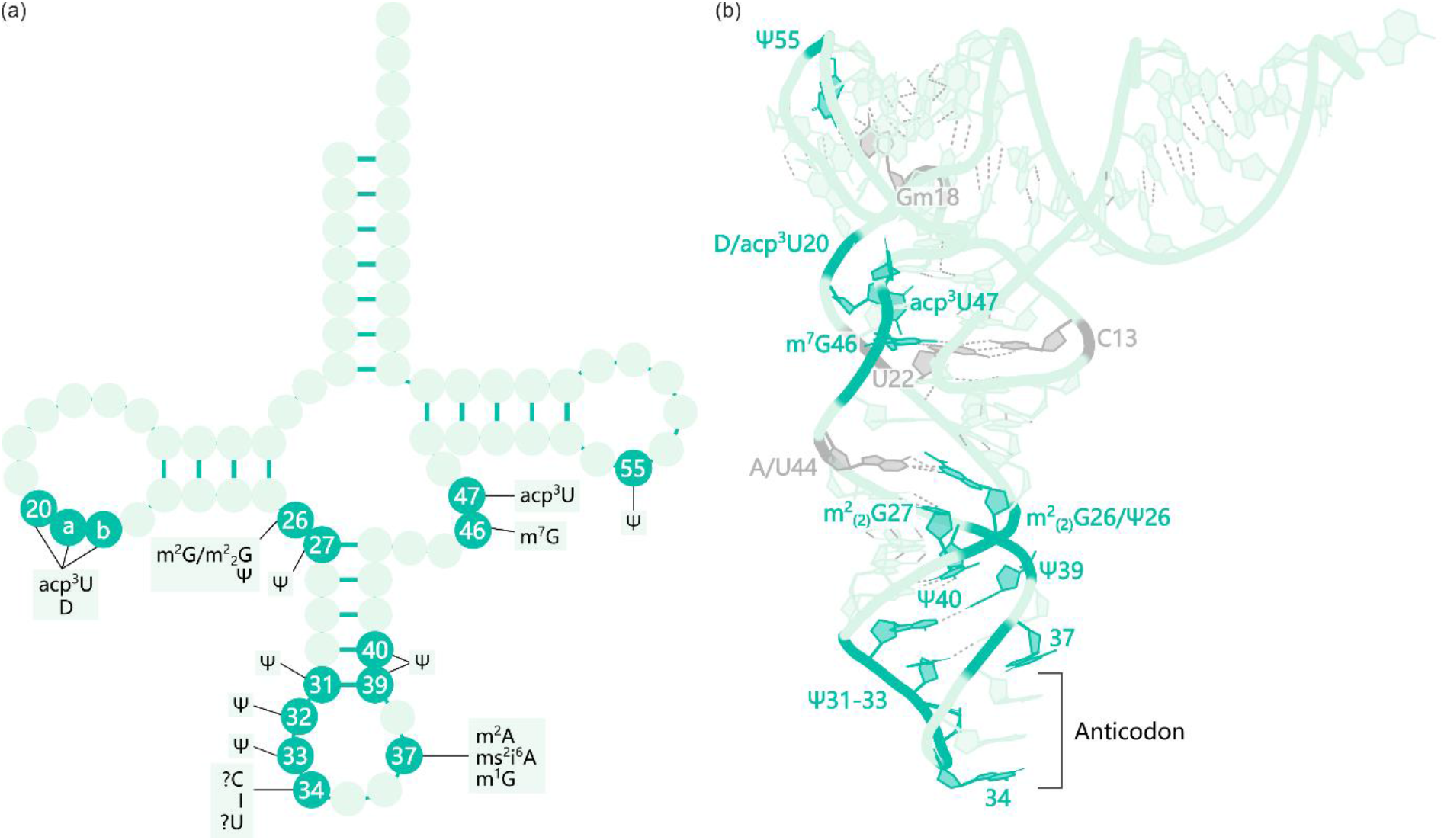
Model of Arabidopsis chloroplast tRNA with modifications. (**a**) The specific locations of these modifications correspond to well-established positions within canonical tRNAs. **(b)** The locations of chemical modifications within the tRNA’s three-dimensional structure. Modified nucleosides are highlighted in dark green. Some of these modified nucleosides form pairs with specific, distantly located nucleosides, which are marked in dark grey. Source: I34, m^2^i^6^A37, m^1^G37, acp^3^U(20,20a,20b), acp^3^U47, m2G/m^2^_2_G26, C34, G17 – this study (tRNA-seq); m^7^G46 – (32); pseudouridine - (100); m^2^A37 - (41). Chemical structures for the presented modifications are provided in Supplementary Figure S1. The tertiary structure of tRNA presented in (**b**) is based on yeast tRNA-Phe^GAA^ (PDB: 1EHZ).

The repertoire of D-arm modifications across species includes dihydrouridine (positions 16, 17, 20, 20a, 20b), pseudouridine (positions 13, 20, 25), acp^3^U (position 20, 20a, 20b), Gm (position 18), m^1^A (position 14, 16), ac^4^C (position 14), and m^3^C (position 20). One highly conserved modification in D-loop, dihydrouridine, serves to improve flexibility by excluding itself from base-stacking interactions and facilitating interactions between adjacent residues in tertiary structure, which are critical for the L-shaped structure (e.g. elbow region). This is also observed in chloroplasts, where at least one dihydrouridine per tRNA is annotated in MODOMICS, confirming the modification’s significance (Supplementary Table 1; (14)). Additionally, many chloroplast tRNAs possess Gm18, which stabilises the tertiary structure by pairing with Ψ55 in bacteria (14). Surprisingly, our sequencing data indicate the presence of acp^3^U at positions 20, 20a, 20b in chloroplast tRNAs. Given that this modification is typically found in eukaryotic tRNAs, its presence in plant organelles was unexpected. However, it has been found in Arabidopsis cytosolic tRNAs (73). The modification diversity at positions 20, 20a and 20b, raises the question of which factors determine whether a specific tRNA nucleoside will be modified into acp^3^U or D. Unlike dihydrouridine, the function of acp^3^U in the D-loop remains elusive. The modification neither affects the nucleoside’s sugar conformation nor participates in Watson-Crick base pairing (53, 74, 75). However, it was suggested that acp^3^U might stabilise the local structure through Mg^2+^ coordination (44, 53). Thus, further investigation of the phenomenon is required.

The genetic code comprises 61 codons that specifically encode amino acids, facilitating the translation of mRNA into polypeptides. The efficiency of translation significantly improves due to the flexibility in genetic information decoding characterised by wobble base pairing. The mechanism allows a single tRNA species to recognise and bind to multiple codons, thereby reducing the number of tRNAs needed to interpret the genetic code (76). The rule relies on noncanonical base pairing of the third codon position in mRNA with the wobble tRNA position (i.e. position 34). As a result, many organisms typically have only 40-50 distinct tRNA species to cover all codons. On the other hand, according to the wobble base pairing, the minimal set of tRNAs consists of 32 species. Since most of plant chloroplast genomes encode only 30 tRNA species, the emergence of the superwobbling concept appears crucial (77). The hypothesis proposes that unmodified U in the first position of the anticodon can pair with all four nucleosides in the third codon position (76). However, the rule contradicts the encoding of multiple distinct amino acids from split codon boxes. Therefore, tRNA modifications at the wobble position are important for accurate and efficient translation (77). Depending on a specific case, they have two general but opposing functions – the increase of codon reading specificity or codon recognition expansion.

Our study provides several examples of chloroplast tRNA modifications at position 34. Firstly, we reconfirmed the presence of I34 in tRNA-Arg^ACG^, which enables it to pair with all four nucleosides in chloroplast (Figure 1h, i; (39, 40)). The same tRNA is modified in prokaryotes, while eukaryotic tRNAs more frequently use the presumptions of I34. Therefore, it indicates that the chloroplast translation system shares similarities with the prokaryotic ancestor. Secondly, we provided information concerning a modification at U34 of tRNA-Leu^UAA^. In *E. coli*, the tRNA species have cmnm^5^s^2^Um34, which occurs only in such tRNA. The modification prevents the misreading of codons ending with U or C, such as UU<u>C</u> encoding phenylalanine (decoded only by one tRNA in *E. coli* and chloroplasts) (78). Although the obtained results did not allow us to analyse RT misincorporation signatures, they certainly proclaim the presence of some modifications at U34 of tRNA-Leu^UAA^ in chloroplasts and *E. coli*. Contrary to the MODOMICS records reporting Um34 found in *Glycine max* and *Phaseolus vulgaris*, our findings suggest that at least in Arabidopsis chloroplast, the tRNA has more complicated modification than ribose methylation, which induces Induro RT hard stops (Supplementary Figure S3a; (14)). The presence of identical modifications at the wobble position in *E. coli* and chloroplast tRNA-Leu^UAA^ remains uncertain; however, our results do not preclude this possibility and thus should be verified. The last detected misincorporation at the wobble position is C34 in tRNA-Ile^CAU^ (Figure 3a-c). In analogy to prokaryotic and archaeal systems, this modification prohibits misreading of methionine codons. Surprisingly, our study suggests the presence of a different modification than bacterial lysidine (Figure 3c). In addition, no ortholog enzyme modifying chloroplast tRNA-Ile^CAU^ has been characterised so far. Thus, the putative novel modification in Arabidopsis chloroplasts awaits its unveiling. Taken together, this study provides novel information about modified tRNAs at the wobble position and suggests the presence of unknown modification types.

Modifications at position 37, 3’-adjacent to the anticodon, improve decoding fidelity. They control the pairing of the codon’s first nucleoside with the anticodon’s third nucleoside (position 36), and maintain an open anticodon loop structure (79). All known tRNAs have a purine (A or G) at position 37, frequently modified in all kingdoms for a specific purpose. For example, m^1^G37 stimulates aminoacylation and prevents +1 frameshifting (80–82). Therefore, the modification installation is crucial for efficient translation as observed in many species’ mutants (83–86). Importantly, two enzymes with distinct origins catalyse the methylation in tRNAs and vary in the recognition mechanism required for their action. Eubacterial TrmD requires a G36G37 sequence motif and structured anticodon and D arms (87–89). In contrast, the action of eukaryotic/archaea Trm5 depends only on the formed tRNA tertiary structure (88, 89). Recently, a putative homolog enzyme of Trm5 (Trm5b, AT4G27340) was identified in Arabidopsis (85). The protein has predicted localisation in chloroplasts and mitochondria, and the homozygous *attrm5b* knock-out mutants are embryo-lethal. Our study indicates that all chloroplast tRNAs with G37 are modified into m^1^G37, including tRNA-Val^GA^**<u>^C^</u>** with C36 (Figure 2d-f, Supplementary Table 1). Therefore, the chloroplast tRNAs are likely ineligible to be methylated by bacterial-type TrmD, and all the findings consistently suggest that the organelles employ the enzyme originating from eukaryotes.

The rest of the tRNAs harbour A37, which is modified most frequently to ms^2^i^6^A, m^6^t^6^A, i^6^A, t^6^A, m^6^A and m^2^A. Threonylcarbamoyladenosine-type (t^6^A, m^6^t^6^A) and isopentyladenosine-type (i^6^A, ms^2^i^6^A) modifications enhance base-stacking to stabilise the codon-anticodon interactions. They also maintain the open structure of the anticodon loop by weakening U33-A37 pairing (Figure 4a, b; (90, 91)). Although our method was insufficient for the detection of most known A37 modifications, it showed the presence of a prokaryotic-type modification – ms^2^i^6^A in most annotated *E. coli* tRNAs (Figure 2a-c, Supplementary Table 2). We also detected the modification in three chloroplast tRNAs using tRNA-seq and supported these observations using LC-MS (Figure 2a-c, Supplementary Tables 1 and 7). The observations are consistent with plastid tRNA MODOMICS records, where also tRNA-Phe^GAA^ and tRNA-Tyr^GUA^ have annotated ms^2^i^6^A37 (annotation of tRNA-Ser^UGA^ is not available). The rest of the annotated A37 modifications include t^6^A, i^6^A, m^6^t^6^A and m^2^A (14). Furthermore, of those modifications that are usually associated with bacteria, m^2^A37 was recently described in chloroplast tRNAs of Arabidopsis (41). The modification strengthens codon-anticodon interactions and accelerates translation elongation. Interestingly, *E. coli* m^2^A methyltransferase RLMN is a dual-target protein modifying rRNA and tRNAs, whereas an enzymatic substrate specialization is observed in plants (41). Collectively, these findings denote a conservation of prokaryotic tRNA modifications at A37 in chloroplast tRNAs.

The variable loop of chloroplast tRNAs shares a modification pattern with prokaryotes. We observe two modifications in this region: m^7^G46 and acp^3^U47. Methylation at G46 is a widespread modification across all life domains. It introduces a positive charge within the tertiary base pair with C13 and U22, thereby stabilising the 3D tRNA core (Figure 4a, b; (92)). Furthermore, its involvement in tRNA modification networks was documented. For instance, m^7^G promotes the formation of acp^3^U in selected *E. coli* tRNAs, and it is associated with Gm, m^5^s^2^U54 and m^1^A58 at high temperatures in *T. thermophilus* (93, 94). The acp^3^U47 is believed to coordinate water or Mg^2+^ in T-arm, which could explain the thermal stability decline observed in mutant strains (44). Although the detected m^7^G46 modifications often do not overlap with acp^3^U47 (Supplementary Table 1), we assume that the presented landscape of m^7^G46 might be incomplete due to the insufficient coverage of some tRNAs in Enroth’s data (32). Therefore, additional verification is required.

Modifications at the T-arm are not highly diverse – each kingdom’s tRNAs contain m^5^U54 and Ψ55 (95). They promote a specific tRNA modification pattern, stabilise the tRNA’s elbow region, and control ribosomal translocation rate (96, 97). In eukaryotes, archaea, thermophilic bacteria, and mitochondria, another residue – m^1^A58 – is important for thermal stability, maturation and global structure of tRNAs (98, 99). Our experiments revealed neither the presence of m^1^A58 in chloroplast tRNAs, nor any other modification. Despite the m^5^U and Ψ detection being inaccessible, the study by Sun *et al.* (100) indicated that Ψ55 is conserved in chloroplast tRNAs. Moreover, all the plastid tRNA records in the MODOMICS database have annotated m^5^U54 and Ψ55. These findings support the conservation of core T-arm modifications in chloroplasts, but their presence needs to be validated.

In conclusion, our study provides a wider view of the chloroplast tRNA modification landscape and reveals undescribed modifications in the organelles so far. Even though the tRNA-seq method has its limitations, it facilitates the detection of acp^3^U (at positions 20, 20a, 20b and 47), m^2^_2_G26, I34, modified C34, cmnm^5^s^2^Um34, ms^2^i^6^37, and m^1^G37. The meta-analysis of already published datasets and literature reports provided information about the presence of m^2^A37, m^7^G46, and Ψ (at positions 26, 27, 31, 32, 33, 39, 40, 55) in Arabidopsis chloroplast tRNAs (Figure 4a, b). Given the nonobvious combination of detected modifications, further exploration of their complete landscape will be a noteworthy area for future research.

### Investigating chloro-ribosome modified nucleosides

In *E. coli*, prior studies have documented 11 modifications in 16S rRNA and 25 modified nucleosides in 23S rRNA, encompassing methylation of the nitrogen bases and ribose, as well as pseudouridylation (14). Leveraging the secondary and cryo-EM structures of the *S. oleracea* chloroplast ribosome, we identified 10 and 22 homologous positions within chloroplast 16S rRNA and 23S rRNA, respectively, which could potentially undergo modifications (Supplementary Table 3, (3)). In some instances, counterpart nucleosides were not found due to the differences in rRNA sequences, including substitutions, deletions, and insertions (Supplementary Figures S6 and S7). Furthermore, we conducted a comprehensive literature review to compile a list of chloroplast rRNA modifications. Research papers provided data on modifications such as pseudouridylation and ribose methylation. For instance, Wu *et al.* (101) detected 5 nucleosides with 2’-O-methylation in the chloro-ribosome, 4 of which are situated at positions homologous to those in prokaryotic ribosomes (two of these positions are conserved across other bacterial species besides *E. coli*). Similarly, Sun *et al.* (100) reported 8 pseudouridines in chloroplast rRNAs with 5 appearing to be plant-specific (see Supplementary Tables 3 and 4, discussed in subsequent sections of this publication).

Subsequently, we conducted an analysis of publicly available Arabidopsis RNA-seq data, with a specific focus on detecting cytosine methylation. Bisulfite conversion stands out as a potent technique for detecting *N*^4^- and 5-methylcytosine (m^4^C and m^5^C) in RNA. This method relies on the selective deamination of unmethylated cytosines (C) to uracils (U) upon bisulfite treatment, while m^5^C and partially m^4^C are protected from conversion (102). We utilised publicly available Arabidopsis data from David *et al.* (31) to detect m^5^C in chloroplast rRNAs. Our analysis reaffirmed the presence of a single m^5^C at position 916 in 16S rRNA (corresponding to m^5^C967 in *E. coli*) and revealed two methylcytosines at positions 1940 and 1977 of 23S rRNA. Notably, m^5^C was not observed at position 1356 in 16S rRNA (the counterpart of *E. coli* m^5^C1407) (refer to Figure 5a and Supplementary Table 3). Our results align with other studies utilising bisulfite conversion on Arabidopsis RNAs, where all the positions were detected as modified (103, 104). However, only Zou *et al.* (104) proposed the existence of a putative plant-specific methylcytosine at positions 897 of 16S rRNA, which was not detected in our analysis due to a low sequencing coverage at this position (approx. 150 reads) (Figure 5a).

**Figure 5.**
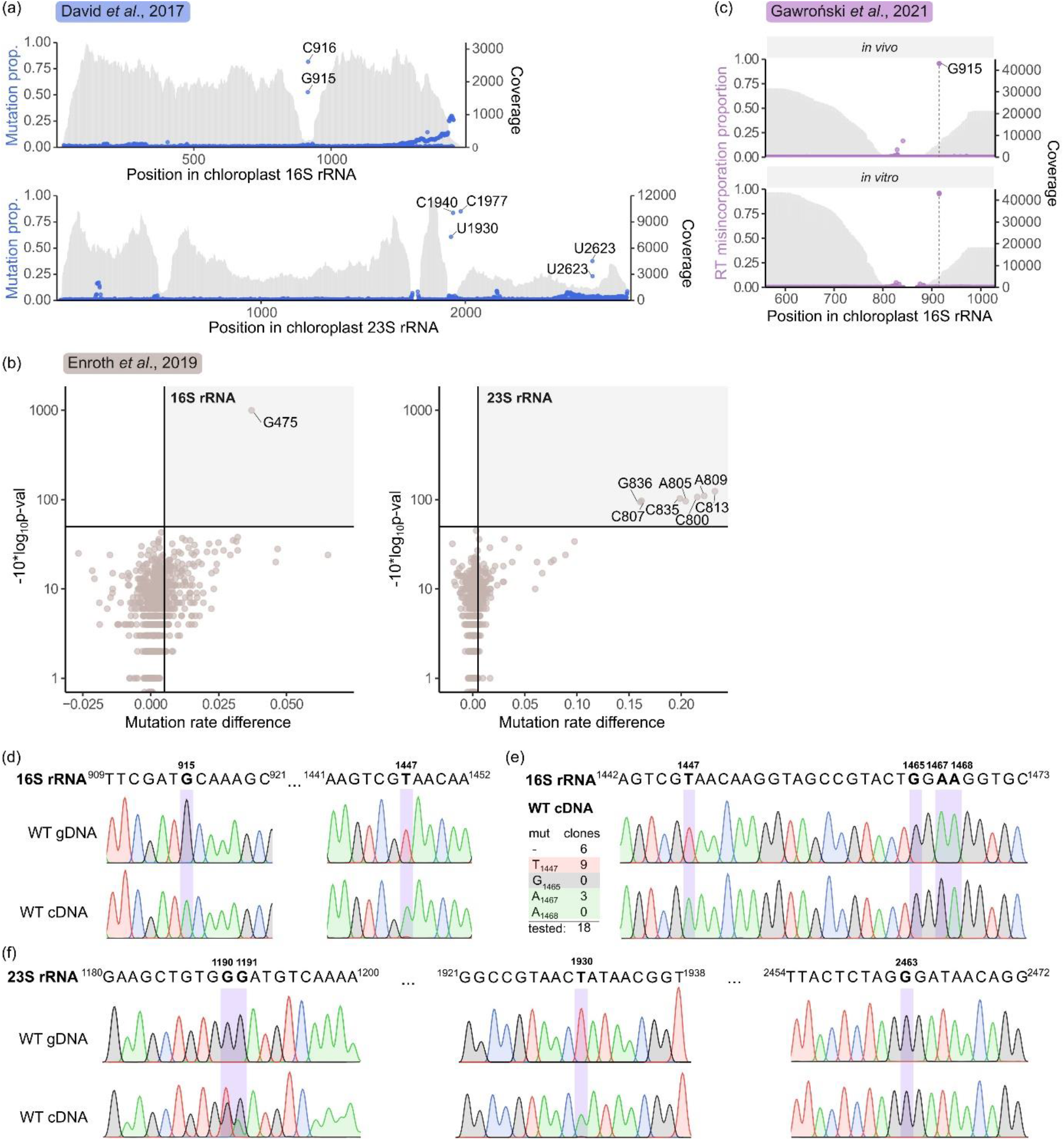
RNA sequencing reveals modified nucleosides in chloro-ribosomes. (**a**) Bisulfite conversion was utilised to detect *N*^4^- and 5-methylcytosine (m^4^C and m^5^C) in 16S rRNA (top) and 23S rRNA (bottom) (31). The fraction of unmodified Cs is shown on the left y-axis and corresponds to m^4^C and/or m^5^C positions indicated on the x-axis. Read coverage is indicated on the right y-axis. Sites with a minimum coverage of 200 reads were analysed. This method also allows for the detection of modifications other than m^4^C and m^5^C that cause reverse transcriptase misincorporations (i.e. G915, U1930, U2622, and U2623). The results represent the average from two replicates. (**b**) Sodium borohydride treatment and mutational profiling were used to detect *N*^7^-methylguanosine (m^7^G) (32). Plots of mutation rate difference (x-axis) versus −10*log(p-val) (y-axis) for 16S rRNA (left) and 23S rRNA (right) are presented. A single m^7^G modification was detected in position 475 of 16S rRNA, with no modifications found in 23S rRNA. (**c**) Mutational profiling was used to detect m^2^G915 in 16S rRNA (30). The fraction of reverse transcriptase (TGIRT-III) misincorporation is shown on the left y-axis and corresponds to positions in 16S rRNA indicated on the x-axis. Read coverage is illustrated on the right y-axis. Sites with a minimum coverage of 500 reads were analysed. The results represent the average from three replicates. (**d-f**) Reverse transcriptase misincorporations at specific sites were analysed using Sanger sequencing of amplified regions using cDNA as a template (bottom). Genomic DNA fragments amplified were used as a negative control (top). (**d**) m^2^G915 and m^3^U1447 were detected in 16S rRNA. (**e**) To detect modifications located close to the 3’ end of 16S rRNA, amplicons were cloned into a vector and 18 clones were sequenced. A table summarising the number of clones with misincorporations is presented on the left, while an example chromatogram of one of the clones is presented on the right. (**f**) Unknown modifications of two Gs at positions 1190 and 1191 (left), m^3^Ψ at position 1930 (middle), and unmodified G2463 corresponding to m^2^G2445 in *E. coli* (right) were analysed in 23S rRNA.

The Enroth *et al.* (32) dataset enabled us to scrutinise two lengthy chloroplast rRNAs (16S rRNA and 23S rRNA), as the authors enriched for longer RNAs. We identified a single m^7^G475 modification in Arabidopsis chloroplast 16S rRNA, corresponding to a modification m^7^G527 in *E. coli* (Figure 5b). However, despite observing enhanced reactivity to NaBH_4_ in certain nucleosides within 23S rRNA, we suspect that they may not represent true m^7^Gs (Figure 5b). This hypothesis is supported by the fact that all highly NaBH_4_-reactive nucleosides cluster within a small region spanning positions 800 to 836 of 23S rRNA. Additionally, most of these nucleosides are not guanosines, and their observed p-values are just above the specified threshold (−10*log_10_(p-value) > 50). In summary, the data indicate the presence of a single m^7^G475 in chloroplast 16S rRNA and raise the possibility that m^7^G may not be present in Arabidopsis chloroplast 23S rRNA, corresponding to *E. coli* m^7^G2069 (Supplementary Table 3).

Reverse transcription can be used to detect RNA modifications occurring on the Watson-Crick face (e.g., m^1^A, m^1^G, m^2^_2_G and m^3^C). This method relies on the reverse transcriptase introducing nucleoside misincorporations into the cDNA at sites of RNA modification. It has proven successful in detecting modified nucleosides in diverse RNA molecules (105–107). Notably, recently developed group II intron RTs, such as TGIRT-III, Marathon-RT, and Induro RT, have demonstrated efficiency in mapping modified nucleosides that lead to premature termination in other RTs or in challenging templates like tRNAs (23, 24, 108). Therefore, we utilised our previous data, employing the high-fidelity TGIRT-III reverse transcriptase, to analyse modifications in positions 560 to 1029 of 16S rRNA (30). Our analysis revealed that G915 exhibits the highest misincorporation rate among analysed samples (Figure 5c). Furthermore, we extended our investigation to the broader region of 16S rRNA utilised in our DMS-seq experiment (29), confirming once again the modification of G915 (Supplementary Figure S5). Importantly, our analysis of m^5^C also showed a high misincorporation rate at G915 (Figure 5a), strongly indicating a modification of this nucleoside on the Watson-Crick face. Consistent with these findings, the *N*^2^-methylguanosine (m^2^G) at position 915 was recently reported in Arabidopsis chloro-ribosomes, catalysed by the RsmD enzyme (109).

Our previously published data, generated with TGIRT-III, omitted the 3’ end of 16S rRNA and the entire 23S rRNA (29, 30). To address this gap, we employed a similar strategy involving Induro RT followed by Sanger sequencing of the chloroplast 16S and 23S rRNA amplicons to expand our understanding of the ribosomal modification landscape. In this experiment, we analysed amplicons obtained from genomic DNA (gDNA) and cDNA templates (Figure 5d, f). The detection of m^2^G915 validated the effectiveness of this strategy (Figure 5d). Additionally, the results revealed a modification at position 1447, corresponding to m^3^U1498 in *E. coli* 16S rRNA (Figure 5d). However, this strategy did not enable us to detect modifications at the very 3’ end of 16S. Therefore, as an additional experiment, we employed circular RT-PCR (cRT-PCR) designed for transcript ends analysis. In this method, circularised RNA is subjected to reverse transcription (SuperScript II RT), resulting in cDNA that serves as a template for amplification, and the PCR products are cloned into vectors for sequencing. Analysis of 18 clones did not reveal any misincorporations at G1465 corresponding to *E. coli* m^2^G1516. Thus, we cannot confirm the conservation of *N*^2^-methylguanosine at this position (Figure 5e). Additionally, we examined neighbouring positions at A1467 and A1468, corresponding to *E. coli* m^6^_2_A1518 and m^6^_2_A1519, respectively. This type of modification disrupts Watson-Crick base pairing, which suggests that it can be detected by our experimental approach. Indeed, we observed misincorporation in that region, but only at A1467 (3 of 18 clones), suggesting the conservation of methylation in one of these residues (Figure 5e). However, we cannot exclude the possibility that the RT used has a lower fidelity towards this type of modification, potentially inhibiting cDNA synthesis. Notably, shortened fragments at 3’ termini are excluded from the analysis in cRT-PCR, with a putative modification at 1468 posing the first obstacle during cDNA synthesis. This hypothesis aligns with results from primer extension assays in Arabidopsis wild-type (WT) plants and *pfc1* mutants lacking the protein methylating these two adenines – KsgA homolog (110). In WT plants, RT stop was observed in the region before A1468, whereas cDNA fragments in *pfc1* were longer and ended at m^3^U1447. Conversely, *Euglena gracilis* chloro-ribosomes exhibit only one dimethylated adenosine – m^6^_2_A1518 (*E. coli* numbering) and yeast mito-ribosomes lack both modifications, suggesting organism-specific variations (111, 112). In light of these results, we confirmed only the presence of modification at A1467, while the conservation of putatively methylated A1468 requires further validation.

Next, we sequenced the 23S rRNA amplicons (divided into three separate fragments). This analysis indicated 3 positions with misincorporations: G1190, G1191 and U1930 (Figure 5f). The first two lack counterparts in *E. coli* and may be specific to plants. In contrast, the presence of misincorporation at U1930 (equivalent to *E. coli* m^3^Ψ1915), suggests the conservation of a modification that affects canonical base pairing. Additionally, we did not observe misincorporations at G2463 – a position corresponding to m^2^G2445 in *E. coli* – indicating that this modification is not conserved in the chloroplast LSU (Figure 5f). Noteworthy, our experimental procedure prevented the analysis of guanine methylation at 1846 (corresponding to *E. coli* m^2^G1835) because it represents the last nucleoside of one of the mature 23S rRNA fragments. In summary, our analysis confirmed 8 prokaryotic-conserved modifications in 16S rRNA and 8 in 23S rRNA, within chloroplast rRNAs (Supplementary Table 3). Additionally, the data unveiled 9 plant-specific chloro-ribosome rRNA modifications (Supplementary Table 4).

### The context of detected chloroplast SSU rRNA modifications

A subset of 16S rRNA modifications playing a pivotal role in the DC was found in chloroplast ribosomes, including m^2^G966, m^5^C967, m^4^Cm1402, m^3^U1498, m^6^_2_A1518, m^6^_2_A1519 (*E. coli* numbering). The modifications are located mainly around the mRNA and tRNA in the P-site, and are highly conserved to maintain a proper geometry of the decoding center. It is not different in chloro-ribosomes, where all the modifications find their counterparts (Figure 6a, b), supporting their crucial role in ribosome functionality.

**Figure 6.**
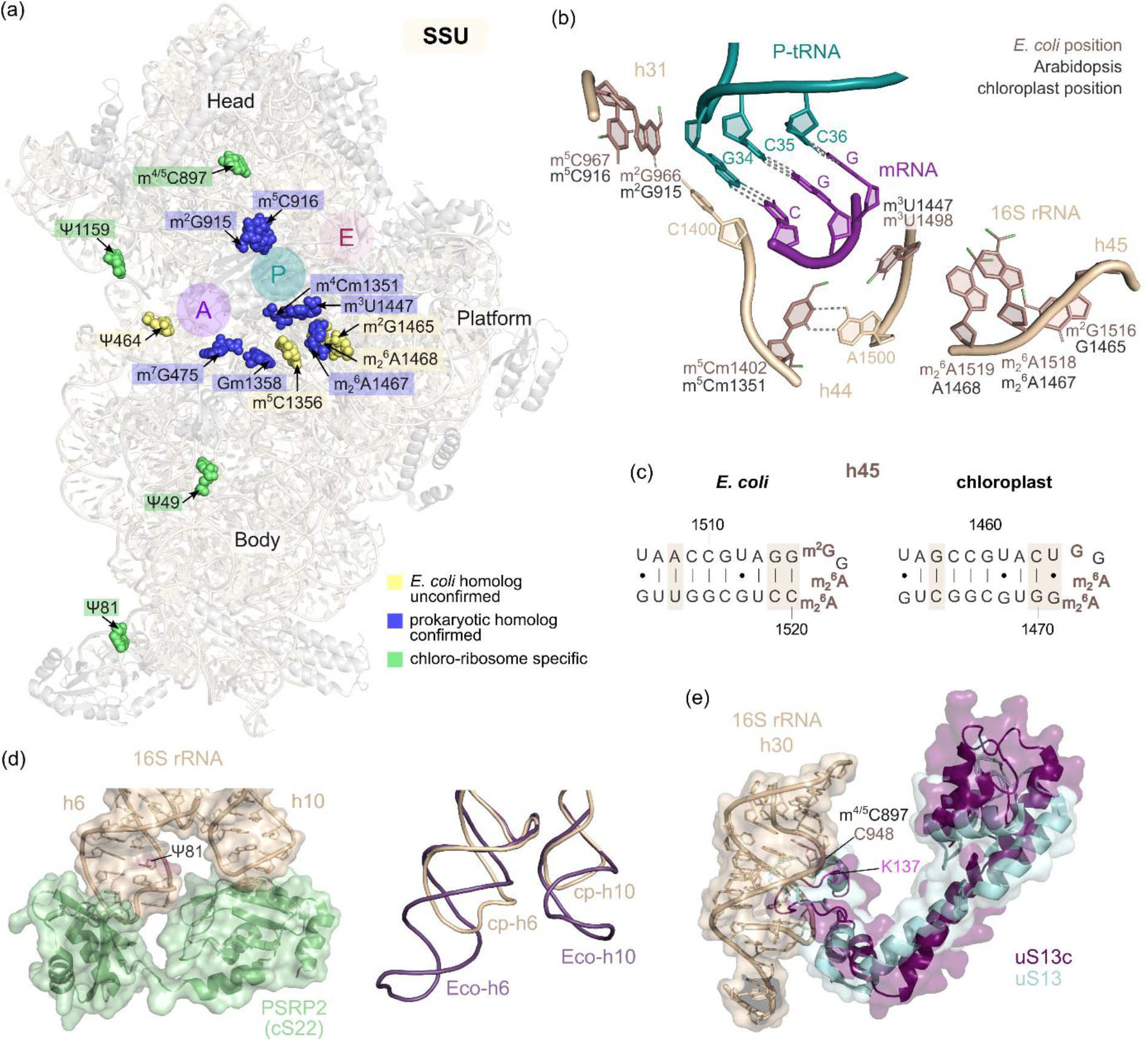
Chloro-specific SSU rRNA modifications likely compensate for changes in structure/composition of SSU and sequence of 16S rRNA. (**a**) The structure of the chloroplast SSU, based on the work of (3) and (6) (PDB: 5MMM – for rRNA and 5X8P – for proteins), with head, platform, and body regions indicated. Modified nucleosides in 16S rRNA homologous to known *E. coli* modifications, confirmed in this study, are marked in blue. Yellow nucleosides represent sites modified in *E. coli* but not observed in chloro-ribosomes, while green nucleosides indicate chloroplast-specific modifications. Most detected modifications are located close to the tRNA binding sites (A, P, and E sites). (**b**) Decoding center architecture with marked nucleosides and specified numbering shown in light brown (*E. coli*) and in grey (Arabidopsis). The tRNA (green) and mRNA (purple) structures (PDB: 6OM6) were superimposed with chloro-ribosome 16S rRNA. (**c**) Secondary structure of helix 45 (h45) in *E. coli* (left) and Arabidopsis (right) 16S rRNA. Sequence differences are highlighted in light brown. Unlike *E. coli*, nucleosides at positions 1465 (corresponding to m^2^G1516) and 1468 (corresponding to m^6^_2_A1519) are not modified in Arabidopsis. (**d**) Plastid-specific ribosomal protein PSRP2 (cS22) binds to helices 6 (h6) and 10 (h10) in close proximity to chloroplast-specific Ψ81 found in h6 (PDB: 5X8R – protein, 5MMJ - rRNA). On the right side, the superimposition of h6 and h10 from *E. coli* (purple, PDB: 6OM6) and chloroplast (light brown, PDB: 5MMJ) is shown. (**e**) The modified C (m^4^C or m^5^C) at position 897 is located in helix 30 (h30, shown in light brown), which interacts with uS15c (purple) (PDB: 5X8R). The chloroplast uS15c was superimposed with bacterial uS13 (light blue) (PDB: 6OM6).

The methylation of nucleosides located in helix 31 at positions 966 and C967 is crucial for recruitment of the initiator tRNA (113). While position 966 (and its equivalents) in most bacteria and chloroplasts contains guanosine, eukaryotes and archaea have uridine at this position. Nevertheless, regardless of the specific nucleoside, a modified nucleoside is always present at this position (114). m^2^G966 establishes a base stacking interaction with m^5^C967, ensuring its appropriate orientation for interaction with the tRNA ribose at position 34 (wobble) in the P site. The function of these modifications results from their interaction with the anticodon loop of tRNA, which is particularly important for the specific binding of tRNA-fMet with the START codon during translation initiation (114, 115). In prokaryotic organisms, RsmD is responsible for incorporating the modification of m^2^G966, while RsmB is responsible for the modification of m^5^C967. RsmB incorporates the modification at the initial stages of SSU assembly by binding to free 16S rRNA, whereas the methylation by RsmD requires prior binding of ribosomal proteins, inducing a conformation change that results in accessible G966 for modification (116). Similarly, our data analysis suggests that chloroplast 16S rRNA undergoes analogous modifications at the equivalents in both of these positions - G915 and C916 (Figure 5a-d; (103, 109, 117). However, only the RsmD homologue (introducing m^2^G915) has been described so far. The analysis of the Arabidopsis mutant defective in RsmD function confirmed its role in 16S rRNA maturation. The *atrsmD* mutants exhibit the accumulation of 16S rRNA precursors, reduced chloroplast-encoded protein levels, and show a cold stress-sensitive phenotype, indicating the role of RsmD in chloroplast translation (109, 117).

Helix 44 is a highly significant region of 16S rRNA due to numerous interactions of its nucleosides with mRNA codons. The second nucleoside of the mRNA codon in the P site interacts with modified 16S rRNA nucleosides - m^4^Cm1402 with the phosphate group, and m^3^U1498 with ribose. Interaction between these modified nucleosides, along with the non-canonical bond between m^4^Cm1402 and A1500, contribute to the stabilization of the local conformation of 16S rRNA in the P site and the recognition of tRNA-fMet (116). Two enzymes are responsible for modification in C1402 - RsmH installs methylation in the nitrogen base and RsmI methylates ribose. In plants, only the CMAL (RsmH homolog) has been identified and it modifies C1351 of chloroplast 16S rRNA (Figure 5a; (104)). Arabidopsis *cmal* mutants show chloroplast translation disorders, along with the accumulation of 16S and 23S rRNA precursors. Unexpectedly, disruptions in the auxin signalling pathway occur, leading to morphological defects in leaves and roots in this mutant (104). Ribose methylation at position 1351 was also detected in the Wu *et al.* (101) data. However, the enzyme responsible for introducing this modification remains uncharacterised (118). Based on a homology search, we speculate that this modification can be catalysed by AT1G45110 (Supplementary Table 3) in Arabidopsis, but further investigation is required. In summary, both modifications are likely conserved at this position in the chloroplast SSU.

RsmE is responsible for modification at m^3^U1498 (119). Although the plant homologue of this gene (putatively AT1G50000) has not yet been functionally characterised, we detected this modification in 16S rRNA as a position with a high misincorporation rate ((118), Figure 5d, e). Moreover, Tokuhisa *et al.* (110) observed RT-stop at this position in primer extension assay in Arabidopsis *pfc1* mutant, suggesting the presence of modified nucleoside.

Among the other conserved modifications within 16S rRNA are the dimethylations of two adenines in helix 45 at positions A1518 and A1519 (*N*^6^,*N*^6^-dimethyladenosine (m^6^_2_ A). The modifications are catalysed by KsgA in a specific mechanism (120). These positions are located in proximity to DC, involving interactions of the P-site tRNA, mRNA and ribosomal LSU. The lack of these modifications induces changes in the geometry of the DC, disrupting the initiation and elongation of translation. The methylations in h45 facilitate its interaction with h44, leading to tertiary structure changes in nucleosides, which directly participate in codon recognition, thus their positional alterations result in reduced accuracy during codon reading and frameshifting (121, 122). The dimethylations occur at a late stage of SSU assembly, thereby facilitating internal quality control. The binding of KsgA prevents the fusion of ribosomal subunits and lasts longer compared to monomethylating enzymes. Consequently, it provides more time to install additional modifications, conformation rearrangement, and factor binding at the final stages of SSU maturation (120, 123). Dissociation of KsgA with Era triggers pre-16S rRNA end processing, releasing SSU nearly prepared for engagement in translation (124). Therefore, KsgA not only introduces modifications crucial for the functional core of the SSU but also prevents premature translation involving immature ribosomal forms. In Arabidopsis, the homolog of KsgA is identified as PFC1 (PALEFACE1), and is likely involved in the methylation of chloroplast 16S rRNA. Arabidopsis *pfc1* mutants subjected to cold stress treatment show a chlorotic phenotype and have impaired chloroplast biogenesis, characteristic of chloroplast translation mutants (110). Intriguingly, bacterial *ksgA* mutants exhibit decelerated growth during cold stress, whereas yeast mutants of the KsgA homologue (DIM1) manifest a lethal phenotype (125). The precise function of the PFC1 protein is unknown. However, the absence of modifications at the 3’ end of the 16S rRNA in *pfc1* mutants suggests PFC1’s potential role in the modification of 16S rRNA at positions corresponding to *E. coli* - A1467 and A1468 (110). Nevertheless, in contrast to A1467, the presence of modification in A1468 remains unconfirmed in plant chloro-ribosomes, especially due to the fact it is not as well conserved as in the proximal adenosine (Figure 5e; (111). Additionally, another modification occurring in the loop of h45 is m^2^G1516, introduced by the RsmJ. This modification does not affect the function of the decoding center (119); therefore, further research is required to examine its function. Notably, we validated neither the presence of the modification in chloroplast SSU nor identified a putative homolog of RsmJ with chloroplast localisation (Figure 5e, Supplementary Table 3). Moreover, we noticed a slight difference in the h45 sequence between *E. coli* and chloroplast 16S rRNA, thus the m^2^G modification in that region might be lost in plants (Figure 6c).

RNA modifications also play a role in antibiotic resistance, which can work in both directions. The antibiotic binding sites and modified nucleosides are often situated near crucial regions for ribosome function. Hence the coincidence of modification appearance and antibiotic targets was questioned (116). In fact, some modifications are necessary for antibiotic action, while the abolition of others facilitates drug susceptibility. For instance, streptomycin binds nearby A-site in SSU and provokes misincorporations in a polypeptide chain. The success of the antibiotic action relies on binding to m^7^G527, as the lack of this modification results in streptomycin resistance (126, 127). The Enroth *et al.* (32) study enabled the detection of m^7^G475 in chloroplast 16S rRNA corresponding with m^7^G527 in *E. coli,* representing the only confirmed *N*^7^-methylguanosine in chloroplast rRNA in the dataset (Figure 5b).

The other antibiotic-dependent modified residues are located in both ribosome subunits – C1409 in h44 of SSU and C1920 in H69 of LSU (*E. coli* numbering). The helices form one of the twelve intersubunit bridges – B2a, which is located just below the A-site. The ribose methylations at these positions are considered crucial for viomycin and capreomycin susceptibility (128, 129). The antibiotics induce intersubunit rotation and stabilise the ribosome conformation in translocation intermediate with tRNAs bound in hybrid states, which results in protein synthesis inhibition (130, 131). Importantly, both modifications appear in *Mycobacterium tuberculosis*, while only Cm1920 in LSU is present in *Thermus thermophilus* and *Campylobacter jejuni*, and neither of them is conserved in *E. coli* (128, 132–135). Interestingly, the expression of *tlyA* (which encodes a protein incorporating these modifications) in *E. coli* also leads to the loss of viomycin and capreomycin resistance (128). In Arabidopsis, Wu *et al.* (101) detected ribose methylation in both counterpart positions – C1358 (SSU) and C1935 (LSU). Furthermore, we unveiled a putative homolog of *M. tuberculosis* TlyA in Arabidopsis (AT3G25470), revealing an evolutional link with the bacteria species. The target of paromomycin – m^5^C1407 is located near mRNA and is important for intersubunit interactions (126). However, we did not detect either the modification in chloroplast equivalent position 1356 in m^5^C profiling data, or RsmF homolog protein (Figure 5a; (31, 103)). Considering the numerous interdependencies between modifications present in the region of 1402-1409 in 16S rRNA, we concluded the absence of m^5^C1356 in chloroplast ribosomes could be somehow compensated by the aforementioned Cm1358 (136). For example, A1408G mutation in bacteria expressing *tlyA* led to improved adaptation to capreomycin, while *N*^1^-methylation of A1408 by CmnU provides resistance to the antibiotic in *Saccharothrix mutabilis* subsp. *Capreolus* (137, 138).

Pseudouridine, a prevalent RNA modification improving structure stability, occurs in prokaryotic 16S rRNA at a single position – 516, catalysed by RsuA. The modification stabilises single RNA conformation, RNA-RNA and RNA-protein interactions due to an extra donor site for hydrogen bond formation, formation of specific water bridges and enhanced base stacking (139–141). Resultantly, pseudouridine increases the thermodynamic stability and rigidity of RNA molecules. In chloroplast SSU, Sun *et al.* (100) identified four SVR1-dependent pseudouridylations in SSU, including Ψ464 (corresponds with *E. coli* Ψ516), and three likely plant-specific Ψ49, Ψ81 and Ψ1159 (Supplementary Tables 3 and 4; (142)). Interestingly, the plant-specific Ψ81 localises in helix 6, which interacts with plastid-specific ribosome protein cS22 - PSRP2 (6). It is tempting to hypothesise the reduction of h6 and h10 lengths and their interaction with PSRP2 is correlated with Ψ81 appearance in plants (Figure 6d).

The other plant-specific pseudouridylation was detected at position 1159, corresponding to U1211 in *E. coli*. This modification is located in h34, where bacteria conserve m^2^G1207, enhancing proper A-site conformation (8). The specific structure of h34 forms a binding pocket for A-site tRNA, where tRNA-N34 directly contacts with 16S rRNA-C1054. It also involves base-pairing C1051 with m^2^G1207 within h34, which is crucial for a functional conformation of the region. Although the modified G1207 does not interact directly with any significant ribosome group, the transversion of G1207 to both pyrimidines caused lethality, due to the base-pairing of ^1054^CA^1055^ with ^1206^GU^1205^ (143, 144). Interestingly, in chloroplast 16S rRNA the base pairing in that region is inversed, resulting in G1000-C1155, and we did not detect the m^2^G1000 modification (Figure S7) suggesting that this modification is absent in chloro-ribosomes. *N*^2^-methylguanosine influences Watson-Crick base pairing, and if present it should be detectable using RNA-seq methods (Figure 5c). However, we cannot rule out the presence of either type of modification or the possibility that the evolution of SVR1-dependent Ψ1159 occurred instead, although their functions may more likely concern A-site ligand binding, for instance (100).

Additionally, Zou *et al.* (104) detected a chloroplast-specific SSU modification - m^4^C or m^5^C at 897 using bisulfite conversion and Sanger sequencing. This nucleoside resides in h30, which interacts with ribosomal protein uS13. Notably, the protein differs slightly from its bacterial homolog primarily due to the N-terminal extension and changes of the overall structure, particularly in the region that interacts with h30 (6). Thus, for example, the closely located K137 of uS13 may interact with modified C897 to compensate for the gaps, but this dependency requires further research (Figure 6e). Conversely, this modification was not detected in C5-methylcytosine transcriptome-wide sequencing (Figure 5a; (31, 103)).

### The context of detected LSU rRNA modifications

In *E. coli*, the most structured rRNA molecule – 23S – contains 24 modified nucleosides, which are well-conserved in many prokaryotic systems, or even universal for all kingdoms. The most significant ones are situated around the A-site, P-site, NPET, and the intersubunit bridge B2a (Figure 7a).

**Figure 7.**
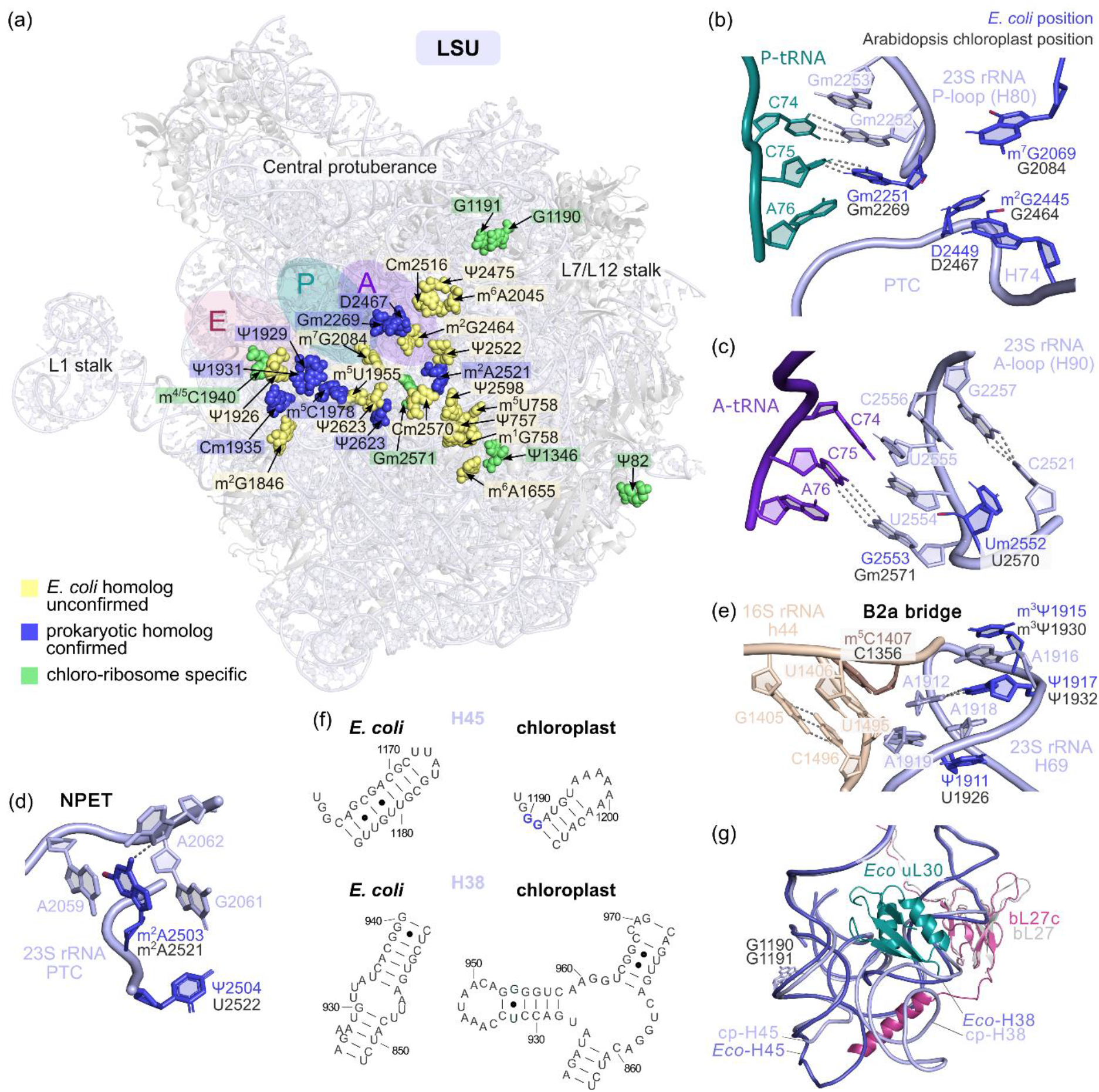
Chloroplast LSU rRNA lacks multiple modifications found in *E. coli* 23S rRNA. (a) The structure of the chloroplast LSU, based on the work of (3) and (6) (PDB: 5MMM – for rRNA and 5X8P – for proteins, 4V4D – for A, P, E-site tRNA locations). Modified nucleosides in 23S rRNA homologous to known *E. coli* modifications, confirmed in this study, are marked in blue. Yellow nucleosides represent sites modified in *E. coli* but not observed in chloro-ribosomes, while green nucleosides indicate chloroplast-specific modifications. Most detected modifications are located close to the tRNA binding sites (A, P, and E sites). (**b**) The structural architecture of the P-loop (H80) interacting with the CCA end of tRNAs (green) in P-site is presented (PDB: 6OM6). The position of modified nucleosides in *E. coli* (blue) and Arabidopsis (grey) are indicated. The modifications in a region of the peptidyl transferase center (PTC) are also shown. (**c**) The structural organisation of the A-loop (H90) interacting with the CCA end of tRNAs (purple) in A-site is presented (PDB: 6OM6, A-site tRNA – 4V42). The position of modified nucleosides in *E. coli* (blue) and Arabidopsis (grey) are indicated. (**d**) The structural organisation of the B2a intersubunit bridge consisting of helix 44 (h44) of 16S rRNA (light brown) and helix 69 (H69) of 23S rRNA (blue) (PDB: 6OM6). The modified nucleosides in *E. coli* (light brown and blue) and their corresponding nucleosides in Arabidopsis (grey) are indicated. The C1356 corresponding to *E. coli* m^5^C1407 is not modified in Arabidopsis. (**e**) The structure and modifications of a region consisting of the nascent peptide exit tunnel (NPET) (PDB: 6OM6). (**f**) Structural organisation of a LSU that contains two chloroplast-specific modifications at positions G1190 and G1191 (PDB: 6OM6, 5X8P). These nucleosides are localised within helix 45 (H45, light blue), which is structurally diverse from its homologous region in *E. coli* (dark blue). The helix 38 (H38), located close to H45, also differs significantly in chloroplasts compared to *E. coli*. The secondary structures of H45 and H38 are presented in panel (**g**).

Two of the most conserved modification sets localise in the P-loop (H80) and A-loop (H90) of 23S rRNA, where ribosome interacts with the CCA-end of tRNAs, controlling their proper positioning. In the P-site, tRNA C74 and C75 pair with 23S rRNA G2252 and Gm2251, respectively. Moreover, the other neighbouring modified nucleosides (m^2^G2445 and D2449) interact with Gm2251, forming a specific structure in the vicinity of the PTC (Figure 7b). Regarding A-site, tRNA C75 pairs with G2553, and Um2552 may improve aminoacyl-tRNA binding (Figure 7c). Despite the similarity of P-site and A-site geometry, the ribose modifications in Gm2251 and Um2552 seem not to be equally important for bacterial viability. Deficiency in Um2552 leads to immature LSU form accumulation and severe growth impairments, which can be suppressed during LSU assembly factor overexpression (i.e. Obg and Der) (145). In turn, there are no phenotypic implications caused by Gm2251 absence in *E. coli* (146). According to Wu *et al.* (101), G2269 (Gm2252 equivalent) is also methylated in chloro-ribosomes. Interestingly, the Um2552 counterpart was not detected as a modified residue, but the neighbouring nucleoside corresponding to *E. coli* G2553 possesses methylated ribose (Gm2571). The situation is not completely specific to plants because the methylation of both Um2552 and G2553 is observed in yeast cytosolic, human cytosolic and mitochondrial ribosomes (147, 148). However, the lack of widely conserved ribose methylation at the 2552 counterpart was not expected and requires additional validation. We can also partially confirm the homology of the other modified residues contacting with Gm2251. In the Sun *et al.* (100) study, Ψ2467 was detected, which corresponds with *E. coli* D2449. However, we did not observe guanine methylation at 2464 (*E. coli* m^2^G2445) in our experiment results or 2084 (*E. coli* m^7^G2069) in Enroth *et al.* (32) (Figure 5b, f). Both modifications are installed by the two-methylase domain enzyme RlmKL, which has no evident homolog in Arabidopsis (118); Supplementary Table 3). Collectively, the data suggest that chloro-ribosome LSU conserves many of the prokaryotic rRNA modifications around the tRNA-binding region but not with absolute fidelity.

Nucleoside modifications within the nascent peptide exit tunnel (NPET) participate in forming a specific trajectory important for nascent protein passage through the ribosome. The NPET contributes to translation regulation by specific interactions of the nascent protein chain and the tunnel wall, which regulate programmed ribosome pausing and translation arrest. The process is regulated by several classes of genes, some of which encode enzymes conferring resistance to macrolide antibiotics (e.g. erythromycin) (149). One of the most conserved modifications – m^2^A2503 interacts with A2059 increasing tunnel wall stability. Moreover, the m^2^A2503 mediates signal transmission between the PTC and NPET, due to its contact with A2062 during one of the ribosome stalling response mechanisms (Figure 7d; (150, 151)). Bacteria deficient in the modified residue exhibit minor growth impairments, but are more sensitive to erythromycin (150). Recently, RLMNL1 incorporating m^2^A2521 (*E. coli* A2503 counterpart) in chloro-ribosomes was identified (41). Although the Arabidopsis *rlmnl1* mutants did not exhibit translation defects in control conditions, they were hypersensitive to paromomycin, which binds SSU in A-site and G418 affecting translation elongation. Moreover, according to Sun *et al.* (100), the neighbouring U2522 is unmodified in contrast to its *E. coli* counterpart Ψ2504 important for PTC inhibitors resistance (Figure 7d; (152)).

The presence of modified residues in the intersubunit bridge region emphasises their role in maintaining ribosome integrity. The abovementioned B2a bridge, composed of h44 and H69, is strengthened by the presence of Ψ1911, m^3^Ψ1915 and Ψ1917 of 23S rRNA. It functions as a part of the decoding center, interacting with tRNAs, release factors and ribosome recycle factors (153). The modified nucleosides in the region interact with other residues via non-canonical Hoogsten base pairing and support subunit associations. In particular, A1912 interacts with Ψ1917 and C1407 of 16S rRNA, while the contact of A1919 with U1406/U1495 of h44 is supported by interactions with A1918 and Ψ1911 (Figure 7e; (153)). In contrast to the wide-conserved pseudouridylations in H69, the modification at a position equivalent to 1915 varies among organisms – there is m^5^U in *T. thermophilus*, and Ψ in archaea and eukaryotes (154). After all, the modification forms a base stacking interaction with A1916, and faces h44, adjusting intersubunit contact for efficient translation in diverse conditions, such as in altered temperature (115). Chloro-ribosomes conserve the modifications corresponding with 1915 and 1917, while the pseudouridylation at 1911 equivalent was not detected. In addition, despite the diversity of modifications at 1915 in diverse organisms, we propound m^3^Ψ conservation in chloroplasts. In contrast to the other known variants in this position, this is the only modification type disturbing Watson-Crick base pairing, which could induce the RT misincorporations that we observed (Figure 5a,f). Interestingly, the bridge contains an additional, plant-specific modification - m^4^C or m^5^C1940, which locates nearby the modified Cm1940 important for antibiotic action (see above).

The two LSU plant-specific pseudouridylations at 1346 (23S) and 82 (4.5S) are likely involved in the distinguishing interactions in chloro-ribosomes. In particular, modification at 4.5S could be important for maintaining the contact of 23S and 4.5S rRNA ends with proteins such as uL3c, bL17c and bL19c (3).

The plant-specific modified guanines at positions 1190 and 1191 are located adjacent to H45, which is shorter compared to its counterpart in *E. coli* (Figure 7f, g). Importantly, the surrounding ribosome region varies significantly in bacteria and chloroplasts in terms of rRNA structure and protein composition. Firstly, the neighbouring H38 contains an additional, chloroplast-specific sequence forming a variable loop within the helix. This adaptation is hypothesised to compensate for the absence of the uL30 homolog in chloro-ribosomes (5). Additionally, the rRNA helices are in close proximity to PSRP6 and bL27c, the chloroplast-distinguished proteins. The bL27c possesses a plant-specific C-terminal extension located between H38 and H45. This protein plays a crucial role in the viability of organisms, both in Arabidopsis and bacteria (155). Therefore, the modified nucleosides specific to chloro-ribosomes could evolve due to the numerous distinctions in this region, making their functional elucidation an attractive area for further research.

In conclusion, we confirmed the presence of 8 prokaryotic homolog modifications and identified 4 plant-specific modified nucleosides in LSU (Supplementary Tables 3 and 4). However, many *E. coli* modifications were not verified based on the sequencing data results. Furthermore, we did not identify putative enzyme homologs incorporating these modifications in LSU using bioinformatic analysis. It is worth noting that, the set of LSU *E. coli* modifications is not universal for all prokaryotes. For instance, the 23S rRNA modification landscape of *Thermus thermophilus* differs significantly from *E. coli* and more closely resembles the one we described for chloro-ribosomes (Supplementary Table 3; (115)). Therefore, the well-described bacterial species may not provide the best reference for the homology of the chloroplast translation machinery.

## SUPPLEMENTARY DATA

Supplementary Data are available at bioRxiv online.

## DATA AVAILABILITY

The code used in this study is available from the corresponding author upon request. The raw NGS fastq (fata) were uploaded to the Sequence Read Archive under BioProject number PRJNA1122064 (https://dataview.ncbi.nlm.nih.gov/object/PRJNA1122064?reviewer=te65rifg2a6io4cdb8h5c1ffrs).

## ACKNOWLEDGEMENTS

The authors wish to thank Dr Nina Sipari at the Viikki Metabolomics Unit, HiLIFE and Faculty of Biological and Environmental Sciences, University of Helsinki, for performing the UPLC-MS runs, and Prof. Damian Gaweł and Anna Bętkowska from the Department of Cell Biology and Immunology, Centre of Postgraduate Medical Education (Warsaw) for *E. coli* str. K-12 substr. MG1665.

## AUTHOR CONTRIBUTIONS

Kinga Gołębiewska: Conceptualization, Formal Analysis, Investigation, Visualization, Writing – original draft, Writing – review & editing. Pavlína Gregorová: Formal Analysis, Investigation, Writing – review & editing. L. Peter Sarin: Conceptualization, Formal Analysis, Supervision, Writing – review & editing. Piotr Gawroński: Conceptualization, Formal Analysis, Supervision, Funding acquisition, Writing – original draft, Writing – review & editing.

## REFERENCES

1. Archibald, J.M. (2015) Genomic perspectives on the birth and spread of plastids. Proc. Natl. Acad. Sci., 112, 10147–10153.

2. Zoschke, R. and Bock, R. (2018) Chloroplast Translation: Structural and Functional Organization, Operational Control, and Regulation. Plant Cell, 30, 745–770.

3. Bieri, P., Leibundgut, M., Saurer, M., Boehringer, D. and Ban, N. (2017) The complete structure of the chloroplast 70S ribosome in complex with translation factor pY. EMBO J., 36, 475–486.

4. Graf, M., Arenz, S., Huter, P., Dönhöfer, A., Nováček, J. and Wilson, D.N. (2016) Cryo-EM structure of the spinach chloroplast ribosome reveals the location of plastid-specific ribosomal proteins and extensions. Nucleic Acids Res., 10.1093/nar/gkw1272.

5. Ahmed, T., Yin, Z. and Bhushan, S. (2016) Cryo-EM structure of the large subunit of the spinach chloroplast ribosome. Sci. Rep., 6, 35793.

6. Ahmed, T., Shi, J. and Bhushan, S. (2017) Unique localization of the plastid-specific ribosomal proteins in the chloroplast ribosome small subunit provides mechanistic insights into the chloroplastic translation. Nucleic Acids Res., 45, 8581–8595.

7. Thiaville, P., Legendre, R., Rojas-Benitez, D., Baudin-Baillieu, A., Hatin, I., Chalancon, G., Glavic, A., Namy, O. and De Crecy-Lagard, V. (2016) Global translational impacts of the loss of the tRNA modification t6A in yeast. Microb. Cell, 3, 29–45.

8. Sergeeva, O.V., Bogdanov, A.A. and Sergiev, P.V. (2015) What do we know about ribosomal RNA methylation in Escherichia coli? Biochimie, 117, 110–118.

9. Fasnacht, M., Gallo, S., Sharma, P., Himmelstoß, M., Limbach, P.A., Willi, J. and Polacek, N. (2022) Dynamic 23S rRNA modification ho5C2501 benefits *Escherichia coli* under oxidative stress. Nucleic Acids Res., 50, 473–489.

10. Lucas, M.C., Pryszcz, L.P., Medina, R., Milenkovic, I., Camacho, N., Marchand, V., Motorin, Y., Ribas De Pouplana, L. and Novoa, E.M. (2024) Quantitative analysis of tRNA abundance and modifications by nanopore RNA sequencing. Nat. Biotechnol., 42, 72–86.

11. Rose, S., Auxilien, S., Havelund, J.F., Kirpekar, F., Huber, H., Grosjean, H. and Douthwaite, S. (2020) The hyperthermophilic partners Nanoarchaeum and Ignicoccus stabilize their tRNA T-loops via different but structurally equivalent modifications. Nucleic Acids Res., 48, 6906–6918.

12. Ohira, T., Minowa, K., Sugiyama, K., Yamashita, S., Sakaguchi, Y., Miyauchi, K., Noguchi, R., Kaneko, A., Orita, I., Fukui, T., et al. (2022) Reversible RNA phosphorylation stabilizes tRNA for cellular thermotolerance. Nature, 605, 372–379.

13. Ohira, T. and Suzuki, T. (2024) Transfer RNA modifications and cellular thermotolerance. Mol. Cell, 84, 94–106.

14. Cappannini, A., Ray, A., Purta, E., Mukherjee, S., Boccaletto, P., Moafinejad, S.N., Lechner, A., Barchet, C., Klaholz, B.P., Stefaniak, F., et al. (2024) MODOMICS: a database of RNA modifications and related information. 2023 update. Nucleic Acids Res., 52, D239–D244.

15. Tuorto, F. and Lyko, F. (2016) Genome recoding by tRNA modifications. Open Biol., 6, 160287.

16. Suzuki, T. (2021) The expanding world of tRNA modifications and their disease relevance. Nat. Rev. Mol. Cell Biol., 22, 375–392.

17. Hernandez-Alias, X., Katanski, C.D., Zhang, W., Assari, M., Watkins, C.P., Schaefer, M.H., Serrano, L. and Pan, T. (2023) Single-read tRNA-seq analysis reveals coordination of tRNA modification and aminoacylation and fragmentation. Nucleic Acids Res., 51, e17–e17.

18. Ashraf, S.S., Sochacka, E., Cain, R., Guenther, R., Malkiewicz, A. and Agris, P.F. (1999) Single atom modification (O-->S) of tRNA confers ribosome binding. RNA N. Y. N, 5, 188–194.

19. Hagelskamp, F., Borland, K., Ramos, J., Hendrick, A.G., Fu, D. and Kellner, S. (2020) Broadly applicable oligonucleotide mass spectrometry for the analysis of RNA writers and erasers in vitro. Nucleic Acids Res., 48, e41–e41.

20. Chan, C.T.Y., Dyavaiah, M., DeMott, M.S., Taghizadeh, K., Dedon, P.C. and Begley, T.J. (2010) A Quantitative Systems Approach Reveals Dynamic Control of tRNA Modifications during Cellular Stress. PLoS Genet., 6, e1001247.

21. Shigematsu, M., Honda, S., Loher, P., Telonis, A.G., Rigoutsos, I. and Kirino, Y. (2017) YAMAT-seq: an efficient method for high-throughput sequencing of mature transfer RNAs. Nucleic Acids Res., 10.1093/nar/gkx005.

22. Pinkard, O., McFarland, S., Sweet, T. and Coller, J. (2020) Quantitative tRNA-sequencing uncovers metazoan tissue-specific tRNA regulation. Nat. Commun., 11, 4104.

23. Behrens, A., Rodschinka, G. and Nedialkova, D.D. (2021) High-resolution quantitative profiling of tRNA abundance and modification status in eukaryotes by mim-tRNAseq. Mol. Cell, 81, 1802–1815.e7.

24. Scheepbouwer, C., Aparicio-Puerta, E., Gomez-Martin, C., Verschueren, H., Van Eijndhoven, M., Wedekind, L.E., Giannoukakos, S., Hijmering, N., Gasparotto, L., Van Der Galien, H.T., et al. (2023) ALL-tRNAseq enables robust tRNA profiling in tissue samples. Genes Dev., 37, 243–257.

25. Sharma, M.R., Wilson, D.N., Datta, P.P., Barat, C., Schluenzen, F., Fucini, P. and Agrawal, R.K. (2007) Cryo-EM study of the spinach chloroplast ribosome reveals the structural and functional roles of plastid-specific ribosomal proteins. Proc. Natl. Acad. Sci., 104, 19315–19320.

26. Perez Boerema, A., Aibara, S., Paul, B., Tobiasson, V., Kimanius, D., Forsberg, B.O., Wallden, K., Lindahl, E. and Amunts, A. (2018) Structure of the chloroplast ribosome with chl-RRF and hibernation-promoting factor. Nat. Plants, 4, 212–217.

27. Rodríguez-Ezpeleta, N., Teijeiro, S., Forget, L., Burger, G. and Lang, B.F. (2009) Construction of cDNA Libraries: Focus on Protists and Fungi. In Parkinson, J. (ed), Expressed Sequence Tags (ESTs), Methods in Molecular Biology. Humana Press, Totowa, NJ, Vol. 533, pp. 33–47.

28. Lampi, M., Gregorova, P., Qasim, M.S., Ahlblad, N.C.V. and Sarin, L.P. (2023) Bacteriophage Infection of the Marine Bacterium Shewanella glacialimarina Induces Dynamic Changes in tRNA Modifications. Microorganisms, 11, 355.

29. Gawroński, P., Pałac, A. and Scharff, L.B. (2020) Secondary Structure of Chloroplast mRNAs In Vivo and In Vitro. Plants, 9, 323.

30. Gawroński, P., Enroth, C., Kindgren, P., Marquardt, S., Karpiński, S., Leister, D., Jensen, P., Vinther, J. and Scharff, L. (2021) Light-Dependent Translation Change of Arabidopsis psbA Correlates with RNA Structure Alterations at the Translation Initiation Region. Cells, 10, 322.

31. David, R., Burgess, A., Parker, B., Li, J., Pulsford, K., Sibbritt, T., Preiss, T. and Searle, I.R. (2017) Transcriptome-Wide Mapping of RNA 5-Methylcytosine in Arabidopsis mRNAs and Noncoding RNAs. Plant Cell, 29, 445–460.

32. Enroth, C., Poulsen, L.D., Iversen, S., Kirpekar, F., Albrechtsen, A. and Vinther, J. (2019) Detection of internal N7-methylguanosine (m7G) RNA modifications by mutational profiling sequencing. Nucleic Acids Res., 47, e126–e126.

33. Kielpinski, L.J., Sidiropoulos, N. and Vinther, J. (2015) Reproducible Analysis of Sequencing-Based RNA Structure Probing Data with User-Friendly Tools. In Methods in Enzymology. Elsevier, Vol. 558, pp. 153–180.

34. Gregorova, P., Sipari, N.H. and Sarin, L.P. (2021) Broad-range RNA modification analysis of complex biological samples using rapid C18-UPLC-MS. RNA Biol., 18, 1382– 1389.

35. Sarin, L.P., Kienast, S.D., Leufken, J., Ross, R.L., Dziergowska, A., Debiec, K., Sochacka, E., Limbach, P.A., Fufezan, C., Drexler, H.C.A., et al. (2018) Nano LC-MS using capillary columns enables accurate quantification of modified ribonucleosides at low femtomol levels. RNA, 24, 1403–1417.

36. De Crécy-Lagard, V. and Jaroch, M. (2021) Functions of Bacterial tRNA Modifications: From Ubiquity to Diversity. Trends Microbiol., 29, 41–53.

37. Gawroński, P., Jensen, P.E., Karpiński, S., Leister, D. and Scharff, L.B. (2018) Pausing of Chloroplast Ribosomes Is Induced by Multiple Features and Is Linked to the Assembly of Photosynthetic Complexes. Plant Physiol., 176, 2557–2569.

38. Yu, N., Jora, M., Solivio, B., Thakur, P., Acevedo-Rocha, C.G., Randau, L., de Crécy-Lagard, V., Addepalli, B. and Limbach, P.A. (2019) tRNA Modification Profiles and Codon-Decoding Strategies in Methanocaldococcus jannaschii. J. Bacteriol., 201, e00690–18.

39. Delannoy, E., Le Ret, M., Faivre-Nitschke, E., Estavillo, G.M., Bergdoll, M., Taylor, N.L., Pogson, B.J., Small, I., Imbault, P. and Gualberto, J.M. (2009) *Arabidopsis* tRNA Adenosine Deaminase Arginine Edits the Wobble Nucleotide of Chloroplast tRNAArg(ACG) and Is Essential for Efficient Chloroplast Translation. Plant Cell, 21, 2058–2071.

40. Karcher, D. and Bock, R. (2009) Identification of the chloroplast adenosine-to-inosine tRNA editing enzyme. RNA, 15, 1251–1257.

41. Duan, H.-C., Zhang, C., Song, P., Yang, J., Wang, Y. and Jia, G. (2024) C2-methyladenosine in tRNA promotes protein translation by facilitating the decoding of tandem m2A-tRNA-dependent codons. Nat. Commun., 15, 1025.

42. Blersch, K.F., Burchert, J.-P., August, S.-C., Welp, L., Neumann, P., Köster, S., Urlaub, H. and Ficner, R. (2021) Structural model of the M7G46 Methyltransferase TrmB in complex with tRNA. RNA Biol., 18, 2466–2479.

43. Ling, J., O’Donoghue, P. and Söll, D. (2015) Genetic code flexibility in microorganisms: novel mechanisms and impact on physiology. Nat. Rev. Microbiol., 13, 707–721.

44. Takakura, M., Ishiguro, K., Akichika, S., Miyauchi, K. and Suzuki, T. (2019) Biogenesis and functions of aminocarboxypropyluridine in tRNA. Nat. Commun., 10, 5542.

45. Suzuki, T. and Numata, T. (2014) Convergent evolution of AUA decoding in bacteria and archaea. RNA Biol., 11, 1586–1596.

46. Alkatib, S., Fleischmann, T.T., Scharff, L.B. and Bock, R. (2012) Evolutionary constraints on the plastid tRNA set decoding methionine and isoleucine. Nucleic Acids Res., 40, 6713–6724.

47. Fages-Lartaud, M. and Hohmann-Marriott, M.F. (2022) Overview of tRNA Modifications in Chloroplasts. Microorganisms, 10, 226.

48. Muramatsu, T., Yokoyama, S., Horie, N., Matsuda, A., Ueda, T., Yamaizumi, Z., Kuchino, Y., Nishimura, S. and Miyazawa, T. (1988) A novel lysine-substituted nucleoside in the first position of the anticodon of minor isoleucine tRNA from Escherichia coli. J. Biol. Chem., 263, 9261–9267.

49. Francis, M.A. and Dudock, B.S. (1982) Nucleotide sequence of a spinach chloroplast isoleucine tRNA. J. Biol. Chem., 257, 11195–11198.

50. Dyubankova, N., Sochacka, E., Kraszewska, K., Nawrot, B., Herdewijn, P. and Lescrinier, E. (2015) Contribution of dihydrouridine in folding of the D-arm in tRNA. Org. Biomol. Chem., 13, 4960–4966.

51. Hendrickson, T.L. (2001) Recognizing the D-loop of transfer RNAs. Proc. Natl. Acad. Sci., 98, 13473–13475.

52. Arrivé, M., Bruggeman, M., Skaltsogiannis, V., Coudray, L., Quan, Y.-F., Schelcher, C., Cognat, V., Hammann, P., Chicher, J., Wolff, P., et al. (2023) A tRNA-modifying enzyme facilitates RNase P activity in Arabidopsis nuclei. Nat. Plants, 9, 2031–2041.

53. Stuart, J.W., Basti, M.M., Smith #, W.S., Forrest #, B., Guenther, R., Sierzputowska-Gracz, H., Nawrot #, B., Malkiewicz, A. and Agris, P.F. (1996) Structure of the Trinucleotide D-acp ^3^ U-A with Coordinated Mg ^2+^ Demonstrates that Modified Nucleosides Contribute to Regional Conformations of RNA. Nucleosides Nucleotides, 15, 1009–1028.

54. Björk, G.R., Durand, J.M.B., Hagervall, T.G., Leipuvien≐, R., Lundgren, H.K., Nilsson, K., Chen, P., Qian, Q. and Urbonavičius, J. (1999) Transfer RNA modification: influence on translational frameshifting and metabolism. FEBS Lett., 452, 47–51.

55. Takeda, H., Hori, H. and Endo, Y. (2002) Identification of Aquifex aeolicus tRNA (m2(2G26) methyltransferase gene. Nucleic Acids Res. Suppl. 2001.

56. Motorin, Y. and Helm, M. (2010) tRNA Stabilization by Modified Nucleotides. Biochemistry, 49, 4934–4944.

57. Kim, S.H., Suddath, F.L., Quigley, G.J., McPherson, A., Sussman, J.L., Wang, A.H.J., Seeman, N.C. and Rich, A. (1974) Three-Dimensional Tertiary Structure of Yeast Phenylalanine Transfer RNA. Science, 185, 435–440.

58. Helm, M., Giegé, R. and Florentz, C. (1999) A Watson−Crick Base-Pair-Disrupting Methyl Group (m ^1^ A9) Is Sufficient for Cloverleaf Folding of Human Mitochondrial tRNA ^Lys^. Biochemistry, 38, 13338–13346.

59. Hori, H. (2019) Regulatory Factors for tRNA Modifications in Extreme-Thermophilic Bacterium Thermus thermophilus. Front. Genet., 10, 204.

60. Nomura, Y., Ohno, S., Nishikawa, K. and Yokogawa, T. (2016) Correlation between the stability of TRNA tertiary structure and the catalytic efficiency of a TRNA -modifying enzyme, archaeal TRNA -guanine transglycosylase. Genes Cells, 21, 41–52.

61. Favre, A., Michelson, A.M. and Yaniv, M. (1971) Photochemistry of 4-thiouridine in Escherichia coli transfer RNA1Val. J. Mol. Biol., 58, 367–379.

62. Thomas, G. and Favre, A. (1975) 4-thiouridine as the target for near-ultraviolet light induced growth delay in Escherichia coli. Biochem. Biophys. Res. Commun., 66, 1454–1461.

63. Pallan, P.S., Kreutz, C., Bosio, S., Micura, R. and Egli, M. (2008) Effects of *N ^2^, N ^2^* - dimethylguanosine on RNA structure and stability: Crystal structure of an RNA duplex with tandem m ^2^ _2_ G:A pairs. RNA, 14, 2125–2135.

64. Sonawane, K.D., Bavi, R.S., Sambhare, S.B. and Fandilolu, P.M. (2016) Comparative Structural Dynamics of tRNAPhe with Respect to Hinge Region Methylated Guanosine: A Computational Approach. Cell Biochem. Biophys., 74, 157–173.

65. Edqvist, J., Grosjean, H. and Stråby, K.B. (1992) Identity elements for N ^2^ -dimethylation of guanosine-26 in yeast tRNAs. Nucleic Acids Res., 20, 6575–6581.

66. Edqvist, J., Blomqvist, K. and Straaby, K. (1994) Structural Elements in Yeast tRNAs Required for Homologous Modification of Guanosine-26 into Dimethylguanosine-26 by the Yeast Trm1 tRNA-Modifying Enzyme. Biochemistry, 33, 9546–9551.

67. Kwong, T.C. and Lane, B.G. (1975) Wheat Embryo Ribonucleates. V. Generation of *N* ^2^ - Dimethylgnanylate When ‘Fully Sequenced’ Homogeneous Species of Transfer RNA are Used as Substrates for Wheat Embryo Methyltransferases. Can. J. Biochem., 53, 690–697.

68. Pegg, A.E. (1974) Sites of methylation of purified transfer ribonucleic acid preparations by enzymes from normal tissues and from tumours induced by dimethylnitrosamine and 1, 2-dimethylhydrazine. Biochem. J., 137, 239–248.

69. Constantinesco, F., Motorin, Y. and Grosjean, H. (1999) Characterisation and Enzymatic Properties of tRNA(guanine 26, N2, N2)-dimethyltransferase (Trm1p) from Pyrococcus furiosus. J. Mol. Biol., 291, 375–392.

70. Awai, T., Kimura, S., Tomikawa, C., Ochi, A., Ihsanawati, Bessho, Y., Yokoyama, S., Ohno, S., Nishikawa, K., Yokogawa, T., et al. (2009) Aquifex aeolicus tRNA (N2, N2-Guanine)-dimethyltransferase (Trm1) Catalyzes Transfer of Methyl Groups Not Only to Guanine 26 but Also to Guanine 27 in tRNA. J. Biol. Chem., 284, 20467–20478.

71. Awai, T., Ochi, A., Ihsanawati, Sengoku, T., Hirata, A., Bessho, Y., Yokoyama, S. and Hori, H. (2011) Substrate tRNA Recognition Mechanism of a Multisite-specific tRNA Methyltransferase, Aquifex aeolicus Trm1, Based on the X-ray Crystal Structure. J. Biol. Chem., 286, 35236–35246.

72. Xiong, Q.-P., Li, J., Li, H., Huang, Z.-X., Dong, H., Wang, E.-D. and Liu, R.-J. (2023) Human TRMT1 catalyzes m2G or m22G formation on tRNAs in a substrate-dependent manner. Sci. China Life Sci., 66, 2295–2309.

73. Funk, H.M., Zhao, R., Thomas, M., Spigelmyer, S.M., Sebree, N.J., Bales, R.O., Burchett, J.B., Mamaril, J.B., Limbach, P.A. and Guy, M.P. (2020) Identification of the enzymes responsible for m2, 2G and acp3U formation on cytosolic tRNA from insects and plants. PLOS ONE, 15, e0242737.

74. Friedman, S., Li, H.J., Nakanishi, K. and Van Lear, G. (1974) 3-(3-Amino-3-carboxypropyl)uridine. Structure of the nucleoside in Escherichia coli transfer ribonucleic acid that reacts with phenoxyacetoxysuccinimide. Biochemistry, 13, 2932– 2937.

75. Friedman, S. (1979) The effect of chemical modification of 3-(3-amino-3-carboxypropyl)uridine on tRNA function. J. Biol. Chem., 254, 7111–7115.

76. Crick, F.H.C. (1966) Codon—anticodon pairing: The wobble hypothesis. J. Mol. Biol., 19, 548–555.

77. Alkatib, S., Scharff, L.B., Rogalski, M., Fleischmann, T.T., Matthes, A., Seeger, S., Schöttler, M.A., Ruf, S. and Bock, R. (2012) The Contributions of Wobbling and Superwobbling to the Reading of the Genetic Code. PLoS Genet., 8, e1003076.

78. Moukadiri, I., Prado, S., Piera, J., Velázquez-Campoy, A., Björk, G.R. and Armengod, M.-E. (2009) Evolutionarily conserved proteins MnmE and GidA catalyze the formation of two methyluridine derivatives at tRNA wobble positions. Nucleic Acids Res., 37, 7177–7193.

79. Seelam Prabhakar, P., Takyi, N.A. and Wetmore, S.D. (2021) Posttranscriptional modifications at the 37th position in the anticodon stem–loop of tRNA: structural insights from MD simulations. RNA, 27, 202–220.

80. Gamper, H.B., Masuda, I., Frenkel-Morgenstern, M. and Hou, Y.-M. (2015) Maintenance of protein synthesis reading frame by EF-P and m1G37-tRNA. Nat. Commun., 6, 7226.

81. Clifton, B.E., Fariz, M.A., Uechi, G.-I. and Laurino, P. (2021) Evolutionary repair reveals an unexpected role of the tRNA modification m1G37 in aminoacylation. Nucleic Acids Res., 49, 12467–12485.

82. Meng, F., Zhou, M., Xiao, Y., Mao, X., Zheng, J., Lin, J., Lin, T., Ye, Z., Cang, X., Fu, Y., et al. (2021) A deafness-associated tRNA mutation caused pleiotropic effects on the m1G37 modification, processing, stability and aminoacylation of tRNAIle and mitochondrial translation. Nucleic Acids Res., 49, 1075–1093.

83. Björk, G.R., Wikström, P.M. and Byström, A.S. (1989) Prevention of Translational Frameshifting by the Modified Nucleoside 1-Methylguanosine. Science, 244, 986– 989.

84. Lee, C., Kramer, G., Graham, D.E. and Appling, D.R. (2007) Yeast Mitochondrial Initiator tRNA Is Methylated at Guanosine 37 by the Trm5-encoded tRNA (Guanine-N1-)-methyltransferase. J. Biol. Chem., 282, 27744–27753.

85. Jin, X., Lv, Z., Gao, J., Zhang, R., Zheng, T., Yin, P., Li, D., Peng, L., Cao, X., Qin, Y., et al. (2019) AtTrm5a catalyses 1-methylguanosine and 1-methylinosine formation on tRNAs and is important for vegetative and reproductive growth in *Arabidopsis thaliana*. Nucleic Acids Res., 47, 883–898.

86. Masuda, I., Hwang, J.-Y., Christian, T., Maharjan, S., Mohammad, F., Gamper, H., Buskirk, A.R. and Hou, Y.-M. (2021) Loss of N1-methylation of G37 in tRNA induces ribosome stalling and reprograms gene expression. eLife, 10, e70619.

87. Redlak, M., Andraos-Selim, C., Giege, R., Florentz, C. and Holmes, W.M. (1997) Interaction of tRNA with tRNA (Guanosine-1)methyltransferase: Binding Specificity Determinants Involve the Dinucleotide G ^36^ pG ^37^ and Tertiary Structure. Biochemistry, 36, 8699–8709.

88. Brulé, H., Elliott, M., Redlak, M., Zehner, Z.E. and Holmes, W.M. (2004) Isolation and Characterization of the Human tRNA-(N ^1^ G37) Methyltransferase (TRM5) and Comparison to the *Escherichia coli* TrmD Protein. Biochemistry, 43, 9243–9255.

89. Christian, T. and Hou, Y.-M. (2007) Distinct Determinants of tRNA Recognition by the TrmD and Trm5 Methyl Transferases. J. Mol. Biol., 373, 623–632.

90. Konevega, A.L., Soboleva, N.G., Makhno, V.I., Semenkov, Y.P., Wintermeyer, W., Rodnina, M.V. and Katunin, V.I. (2004) Purine bases at position 37 of tRNA stabilize codon–anticodon interaction in the ribosomal A site by stacking and Mg ^2+^ -dependent interactions. RNA, 10, 90–101.

91. Smith, T.J., Giles, R.N. and Koutmou, K.S. (2024) Anticodon stem-loop tRNA modifications influence codon decoding and frame maintenance during translation. Semin. Cell Dev. Biol., 154, 105–113.

92. Hoburg, A., Aschhoff, H.J., Kersten, H., Manderschied, U. and Gassen, H.G. (1979) Function of Modified Nucleosides 7-Methylguanosine, Ribothymidine, and 2-Thiomethyl-*N* ^6^ -(Isopentenyl)adenosine in Procaryotic Transfer Ribonucleic Acid. J. Bacteriol., 140, 408–414.

93. Meyer, B., Immer, C., Kaiser, S., Sharma, S., Yang, J., Watzinger, P., Weiß, L., Kotter, A., Helm, M., Seitz, H.-M., et al. (2020) Identification of the 3-amino-3-carboxypropyl (acp) transferase enzyme responsible for acp3U formation at position 47 in Escherichia coli tRNAs. Nucleic Acids Res., 48, 1435–1450.

94. Droogmans, L. (2003) Cloning and characterization of tRNA (m1A58) methyltransferase (TrmI) from Thermus thermophilus HB27, a protein required for cell growth at extreme temperatures. Nucleic Acids Res., 31, 2148–2156.

95. Roovers, M., Droogmans, L. and Grosjean, H. (2021) Post-Transcriptional Modifications of Conserved Nucleotides in the T-Loop of tRNA: A Tale of Functional Convergent Evolution. Genes, 12, 140.

96. Pan, D., Kirillov, S., Zhang, C.-M., Hou, Y.-M. and Cooperman, B.S. (2006) Rapid ribosomal translocation depends on the conserved 18-55 base pair in P-site transfer RNA. Nat. Struct. Mol. Biol., 13, 354–359.

97. Jones, J.D., Franco, M.K., Tardu, M., Smith, T.J., Snyder, L.R., Eyler, D.E., Polikanov, Y., Kennedy, R.T., Niederer, R.O. and Koutmou, K.S. (2023) Conserved 5-methyluridine tRNA modification modulates ribosome translocation. 10.1101/2023.11.12.566704.

98. Kuratani, M., Yanagisawa, T., Ishii, R., Matsuno, M., Si, S.-Y., Katsura, K., Ushikoshi-Nakayama, R., Shibata, R., Shirouzu, M., Bessho, Y., et al. (2014) Crystal structure of tRNA m1A58 methyltransferase TrmI from Aquifex aeolicus in complex with S-adenosyl-l-methionine. J. Struct. Funct. Genomics, 15, 173–180.

99. Yared, M.-J., Yoluç, Y., Catala, M., Tisné, C., Kaiser, S. and Barraud, P. (2023) Different modification pathways for m1A58 incorporation in yeast elongator and initiator tRNAs. Nucleic Acids Res., 51, 10653–10667.

100. Sun, L., Xu, Y., Bai, S., Bai, X., Zhu, H., Dong, H., Wang, W., Zhu, X., Hao, F. and Song, C.-P. (2019) Transcriptome-wide analysis of pseudouridylation of mRNA and non-coding RNAs in Arabidopsis. J. Exp. Bot., 70, 5089–5600.

101. Wu, S., Wang, Y., Wang, J., Li, X., Li, J. and Ye, K. (2021) Profiling of RNA ribose methylation in *Arabidopsis thaliana*. Nucleic Acids Res., 49, 4104–4119.

102. Dai, Q., Ye, C., Irkliyenko, I., Wang, Y., Sun, H.-L., Gao, Y., Liu, Y., Beadell, A., Perea, J., Goel, A., et al. (2024) Ultrafast bisulfite sequencing detection of 5-methylcytosine in DNA and RNA. Nat. Biotechnol., 10.1038/s41587-023-02034-w.

103. Burgess, A.L., David, R. and Searle, I.R. (2015) Conservation of tRNA and rRNA 5-methylcytosine in the kingdom Plantae. BMC Plant Biol., 15, 199.

104. Zou, M., Mu, Y., Chai, X., Ouyang, M., Yu, L.-J., Zhang, L., Meurer, J. and Chi, W. (2020) The critical function of the plastid rRNA methyltransferase, CMAL, in ribosome biogenesis and plant development. Nucleic Acids Res., 48, 3195–3210.

105. Ebhardt, H.A., Tsang, H.H., Dai, D.C., Liu, Y., Bostan, B. and Fahlman, R.P. (2009) Meta-analysis of small RNA-sequencing errors reveals ubiquitous post-transcriptional RNA modifications. Nucleic Acids Res., 37, 2461–2470.

106. Ryvkin, P., Leung, Y.Y., Silverman, I.M., Childress, M., Valladares, O., Dragomir, I., Gregory, B.D. and Wang, L.-S. (2013) HAMR: high-throughput annotation of modified ribonucleotides. RNA, 19, 1684–1692.

107. Hauenschild, R., Tserovski, L., Schmid, K., Thüring, K., Winz, M.-L., Sharma, S., Entian, K.-D., Wacheul, L., Lafontaine, D.L.J., Anderson, J., et al. (2015) The reverse transcription signature of *N*-1-methyladenosine in RNA-Seq is sequence dependent. Nucleic Acids Res., 10.1093/nar/gkv895.

108. Nakano, Y., Gamper, H., McGuigan, H., Maharjan, S., Sun, Z., Krishnan, K., Yigit, E., Li, N.-S., Piccirilli, J.A., Kleiner, R., et al. (2023) Genome-Wide Profiling of tRNA Using an Unexplored Reverse Transcriptase with High Processivity. 10.1101/2023.12.09.569604.

109. Ngoc, L.N.T., Park, S.J., Cai, J., Huong, T.T., Lee, K. and Kang, H. (2021) RsmD, a Chloroplast rRNA m2G Methyltransferase, Plays a Role in Cold Stress Tolerance by Possibly Affecting Chloroplast Translation in *Arabidopsis*. Plant Cell Physiol., 62, 948–958.

110. Tokuhisa, J.G., Vijayan, P., Feldmann, K.A. and Browse, J.A. (1998) Chloroplast Development at Low Temperatures Requires a Homolog of *DIM1*, a Yeast Gene Encoding the 18S rRNA Dimethylase. Plant Cell, 10, 699–711.

111. Van Buul, C.P.J.J., Hamersma, M., Visser, W. and Van Knippenberg, P.H. (1984) Partial methylation of two adjacent adenosines in ribosomes from *Euglena gracilis* chloroplasts suggests evolutionary loss of an intermediate stage in the methyl-transfer reaction. Nucleic Acids Res., 12, 9205–9208.

112. Klootwijk, J., Klein, I. and Grivell, L.A. (1975) Minimal post-transcriptional modification of yeast mitochondrial ribosomal RNA. J. Mol. Biol., 97, 337–350.

113. Burakovsky, D.E., Prokhorova, I.V., Sergiev, P.V., Milón, P., Sergeeva, O.V., Bogdanov, A.A., Rodnina, M.V. and Dontsova, O.A. (2012) Impact of methylations of m2G966/m5C967 in 16S rRNA on bacterial fitness and translation initiation. Nucleic Acids Res., 40, 7885–7895.

114. Sergiev, P.V., Aleksashin, N.A., Chugunova, A.A., Polikanov, Y.S. and Dontsova, O.A. (2018) Structural and evolutionary insights into ribosomal RNA methylation. Nat. Chem. Biol., 14, 226–235.

115. Polikanov, Y.S., Melnikov, S.V., Söll, D. and Steitz, T.A. (2015) Structural insights into the role of rRNA modifications in protein synthesis and ribosome assembly. Nat. Struct. Mol. Biol., 22, 342–344.

116. Sergiev, P.V., Golovina, A.Y., Prokhorova, I.V., Sergeeva, O.V., Osterman, I.A., Nesterchuk, M.V., Burakovsky, D.E., Bogdanov, A.A. and Dontsova, O.A. (2011) Modifications of ribosomal RNA: From enzymes to function. In Rodnina, M.V., Wintermeyer, W., Green, R. (eds), Ribosomes. Springer Vienna, Vienna, pp. 97–110.

117. Liu, K., Lee, K.P., Duan, J., Kim, E.Y., Singh, R.M., Di, M., Meng, Z. and Kim, C. (2023) Cooperative role of ATRSMD and ATRIMM proteins in modification and maturation of 16S RRNA in plastids. Plant J., 114, 310–324.

118. Manduzio, S. and Kang, H. (2021) RNA methylation in chloroplasts or mitochondria in plants. RNA Biol., 18, 2127–2135.

119. Basturea, G.N., Rudd, K.E. and Deutscher, M.P. (2006) Identification and characterization of RsmE, the founding member of a new RNA base methyltransferase family. RNA, 12, 426–434.

120. Stephan, N.C., Ries, A.B., Boehringer, D. and Ban, N. (2021) Structural basis of successive adenosine modifications by the conserved ribosomal methyltransferase KsgA. Nucleic Acids Res., 49, 6389–6398.

121. O’Connor, M. (1997) Decoding fidelity at the ribosomal A and P sites: influence of mutations in three different regions of the decoding domain in 16S rRNA. Nucleic Acids Res., 25, 1185–1193.

122. Demirci, H., Murphy, F., Belardinelli, R., Kelley, A.C., Ramakrishnan, V., Gregory, S.T., Dahlberg, A.E. and Jogl, G. (2010) Modification of 16S ribosomal RNA by the KsgA methyltransferase restructures the 30S subunit to optimize ribosome function. RNA, 16, 2319–2324.

123. Sun, J., Kinman, L.F., Jahagirdar, D., Ortega, J. and Davis, J.H. (2023) KsgA facilitates ribosomal small subunit maturation by proofreading a key structural lesion. Nat. Struct. Mol. Biol., 30, 1468–1480.

124. Maksimova, E., Kravchenko, O., Korepanov, A. and Stolboushkina, E. (2022) Protein Assistants of Small Ribosomal Subunit Biogenesis in Bacteria. Microorganisms, 10, 747.

125. Lafontaine, D., Vandenhaute, J. and Tollervey, D. (1995) The 18S rRNA dimethylase Dim1p is required for pre-ribosomal RNA processing in yeast. Genes Dev., 9, 2470– 2481.

126. Carter, A.P., Clemons, W.M., Brodersen, D.E., Morgan-Warren, R.J., Wimberly, B.T. and Ramakrishnan, V. (2000) Functional insights from the structure of the 30S ribosomal subunit and its interactions with antibiotics. Nature, 407, 340–348.

127. Okamoto, S., Tamaru, A., Nakajima, C., Nishimura, K., Tanaka, Y., Tokuyama, S., Suzuki, Y. and Ochi, K. (2007) Loss of a conserved 7-methylguanosine modification in 16S rRNA confers low-level streptomycin resistance in bacteria. Mol. Microbiol., 63, 1096–1106.

128. Johansen, S.K., Maus, C.E., Plikaytis, B.B. and Douthwaite, S. (2006) Capreomycin Binds across the Ribosomal Subunit Interface Using tlyA-Encoded 2′-O-Methylations in 16S and 23S rRNAs. Mol. Cell, 23, 173–182.

129. Maus, C.E., Plikaytis, B.B. and Shinnick, T.M. (2005) Mutation of *tlyA* Confers Capreomycin Resistance in *Mycobacterium tuberculosis*. Antimicrob. Agents Chemother., 49, 571–577.

130. Ermolenko, D.N., Spiegel, P.C., Majumdar, Z.K., Hickerson, R.P., Clegg, R.M. and Noller, H.F. (2007) The antibiotic viomycin traps the ribosome in an intermediate state of translocation. Nat. Struct. Mol. Biol., 14, 493–497.

131. Zhang, L., Wang, Y.-H., Zhang, X., Lancaster, L., Zhou, J. and Noller, H.F. (2020) The structural basis for inhibition of ribosomal translocation by viomycin. Proc. Natl. Acad. Sci., 117, 10271–10277.

132. Guymon, R., Pomerantz, S.C., Crain, P.F. and McCloskey, J.A. (2006) Influence of Phylogeny on Posttranscriptional Modification of rRNA in Thermophilic Prokaryotes: The Complete Modification Map of 16S rRNA of *Thermus thermophilus*. Biochemistry, 45, 4888–4899.

133. Mengel-Jørgensen, J., Jensen, S.S., Rasmussen, A., Poehlsgaard, J., Iversen, J.J.L. and Kirpekar, F. (2006) Modifications in Thermus thermophilus 23 S Ribosomal RNA Are Centered in Regions of RNA-RNA Contact. J. Biol. Chem., 281, 22108–22117.

134. Monshupanee, T., Johansen, S.K., Dahlberg, A.E. and Douthwaite, S. (2012) Capreomycin susceptibility is increased by TlyA-directed 2′-*O* -methylation on both ribosomal subunits. Mol. Microbiol., 85, 1194–1203.

135. Sałamaszyńska-Guz, A., Rose, S., Lykkebo, C.A., Taciak, B., Bącal, P., Uśpieński, T. and Douthwaite, S. (2018) Biofilm Formation and Motility Are Promoted by Cj0588-Directed Methylation of rRNA in Campylobacter jejuni. Front. Cell. Infect. Microbiol., 7, 533.

136. Osterman, I.A., Dontsova, O.A. and Sergiev, P.V. (2020) rRNA Methylation and Antibiotic Resistance. Biochem. Mosc., 85, 1335–1349.

137. Freihofer, P., Akbergenov, R., Teo, Y., Juskeviciene, R., Andersson, D.I. and Böttger, E.C. (2016) Nonmutational compensation of the fitness cost of antibiotic resistance in mycobacteria by overexpression of *tlyA* rRNA methylase. RNA, 22, 1836–1843.

138. Felnagle, E.A., Rondon, M.R., Berti, A.D., Crosby, H.A. and Thomas, M.G. (2007) Identification of the Biosynthetic Gene Cluster and an Additional Gene for Resistance to the Antituberculosis Drug Capreomycin. Appl. Environ. Microbiol., 73, 4162–4170.

139. Davis, D.R. (1995) Stabilization of RNA stacking by pseudouridine. Nucleic Acids Res., 23, 5020–5026.

140. Deb, I., Popenda, Ł., Sarzyńska, J., Małgowska, M., Lahiri, A., Gdaniec, Z. and Kierzek, R. (2019) Computational and NMR studies of RNA duplexes with an internal pseudouridine-adenosine base pair. Sci. Rep., 9, 16278.

141. Borchardt, E.K., Martinez, N.M. and Gilbert, W.V. (2020) Regulation and Function of RNA Pseudouridylation in Human Cells. Annu. Rev. Genet., 54, 309–336.

142. Yu, F., Liu, X., Alsheikh, M., Park, S. and Rodermel, S. (2008) Mutations in *SUPPRESSOR OF VARIEGATION1*, a Factor Required for Normal Chloroplast Translation, Suppress *var2* -Mediated Leaf Variegation in *Arabidopsis*. Plant Cell, 20, 1786–1804.

143. Jemiolo, D.K., Taurence, J.S. and Giese, S. (1991) Mutations in 16S rRNA in *Escherichia coli* at methyl-modified sites: G966, C967, and G1207. Nucleic Acids Res., 19, 4259–4265.

144. Sergiev, P.V., Bogdanov, A.A. and Dontsova, O.A. (2007) Ribosomal RNA guanine-(N2)-methyltransferases and their targets. Nucleic Acids Res., 35, 2295–2301.

145. Lövgren, J.M. and Wikström, P.M. (2001) The *rlmB* Gene Is Essential for Formation of Gm2251 in 23S rRNA but Not for Ribosome Maturation in *Escherichia coli*. J. Bacteriol., 183, 6957–6960.

146. Caldas, T., Binet, E., Bouloc, P. and Richarme, G. (2000) Translational Defects of Escherichia coli Mutants Deficient in the Um2552 23S Ribosomal RNA Methyltransferase RrmJ/FTSJ. Biochem. Biophys. Res. Commun., 271, 714–718.

147. Lapeyre, B. and Purushothaman, S.K. (2004) Spb1p-Directed Formation of Gm2922 in the Ribosome Catalytic Center Occurs at a Late Processing Stage. Mol. Cell, 16, 663– 669.

148. Lee, K.-W. and Bogenhagen, D.F. (2014) Assignment of 2′-O-Methyltransferases to Modification Sites on the Mammalian Mitochondrial Large Subunit 16 S Ribosomal RNA (rRNA). J. Biol. Chem., 289, 24936–24942.

149. Ramu, H., Mankin, A. and Vazquez-Laslop, N. (2009) Programmed drug-dependent ribosome stalling. Mol. Microbiol., 71, 811–824.

150. Vázquez-Laslop, N., Ramu, H., Klepacki, D., Kannan, K. and Mankin, A.S. (2010) The key function of a conserved and modified rRNA residue in the ribosomal response to the nascent peptide. EMBO J., 29, 3108–3117.

151. Koch, M., Willi, J., Pradère, U., Hall, J. and Polacek, N. (2017) Critical 23S rRNA interactions for macrolide-dependent ribosome stalling on the ErmCL nascent peptide chain. Nucleic Acids Res., 45, 6717–6728.

152. Toh, S.-M. and Mankin, A.S. (2008) An Indigenous Posttranscriptional Modification in the Ribosomal Peptidyl Transferase Center Confers Resistance to an Array of Protein Synthesis Inhibitors. J. Mol. Biol., 380, 593–597.

153. Sakakibara, Y. and Chow, C.S. (2012) Role of Pseudouridine in Structural Rearrangements of Helix 69 During Bacterial Ribosome Assembly. ACS Chem. Biol., 7, 871–878.

154. Piekna-Przybylska, D., Decatur, W.A. and Fournier, M.J. (2007) The 3D rRNA modification maps database: with interactive tools for ribosome analysis. Nucleic Acids Res., 36, D178–D183.

155. Romani, I., Tadini, L., Rossi, F., Masiero, S., Pribil, M., Jahns, P., Kater, M., Leister, D. and Pesaresi, P. (2012) Versatile roles of Arabidopsis plastid ribosomal proteins in plant growth and development. Plant J., 72, 922–934.

156. Edqvist, J., Ståby, K.B. and Grosjean, H. (1995) Enzymatic formation of N2, N2-dimethylguanosine in eukaryotic tRNA: Importance of the tRNA architecture. Biochimie, 77, 54–61.

