## Supplementary figures for "Deciphering the RNA Modification Landscape in Arabidopsis Chloroplast tRNAs and rRNAs Reveals a Blend of Ancestral and Acquired Characteristics"

### MANUSCRIPT TITLE

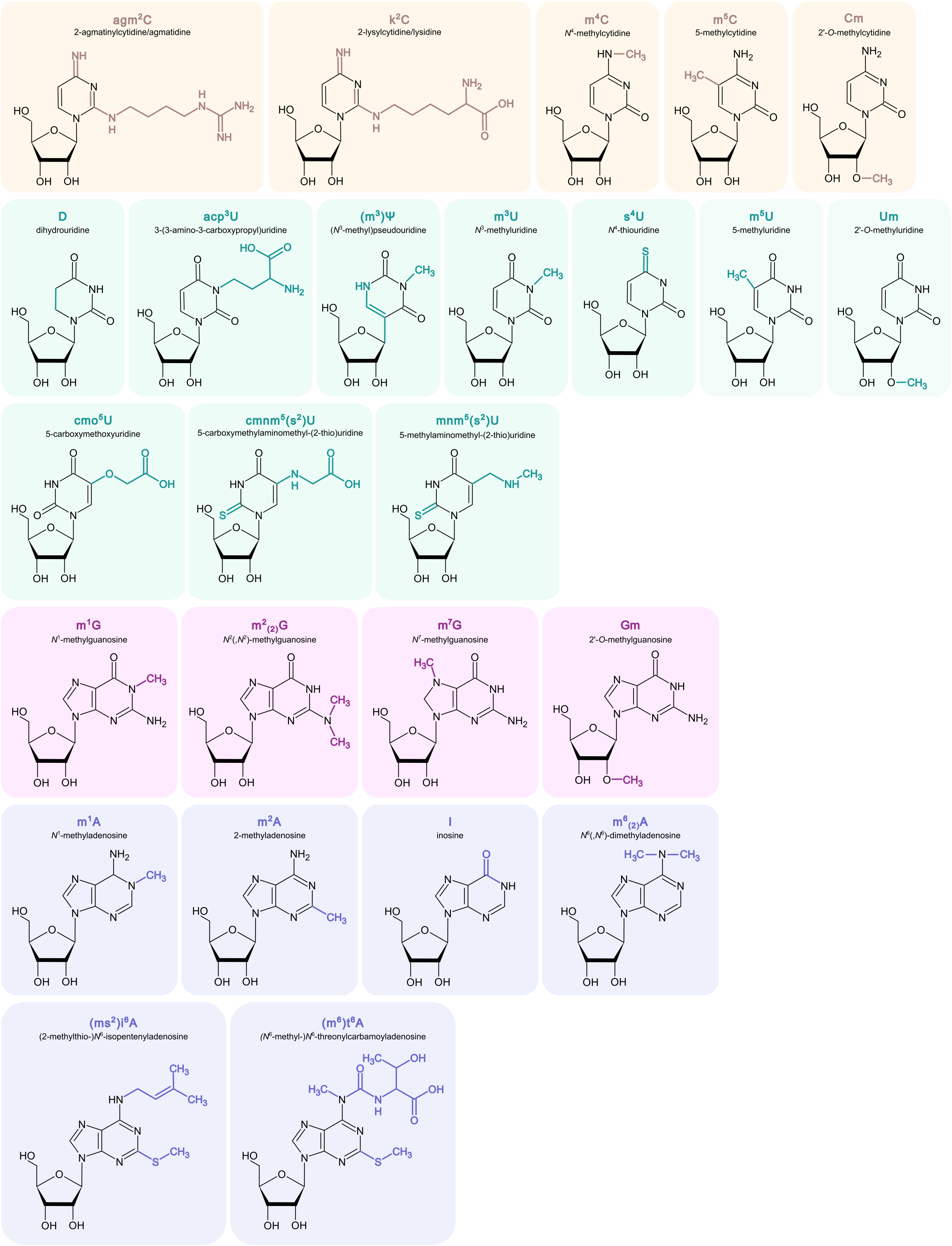


**Figure S1. Chemical structures of modified nucleotides in tRNAs and rRNAs.** The derivatives of cytidine, uridine, guanosine, and adenosine are presented on light orange, green, purple, and blue backgrounds, respectively.

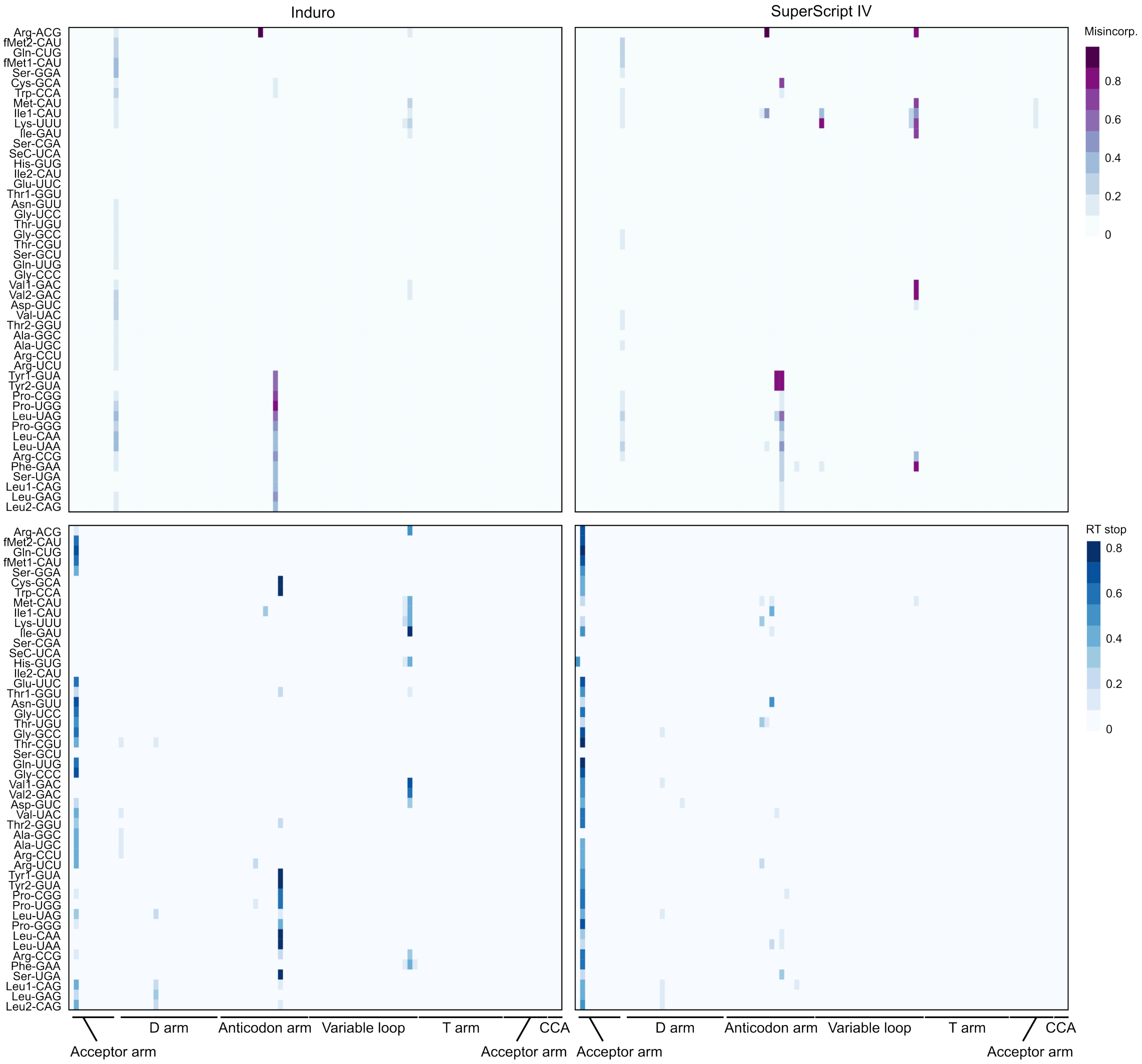


**Figure S2. RT Misincorporation and RT stop heat maps – *E. coli* tRNAs.** Heatmaps illustrating the frequency of RT misincorporations (top) and RT stops (bottom) for Induro (left) and SuperScript IV (right) RTs across *E. coli* tRNAs.

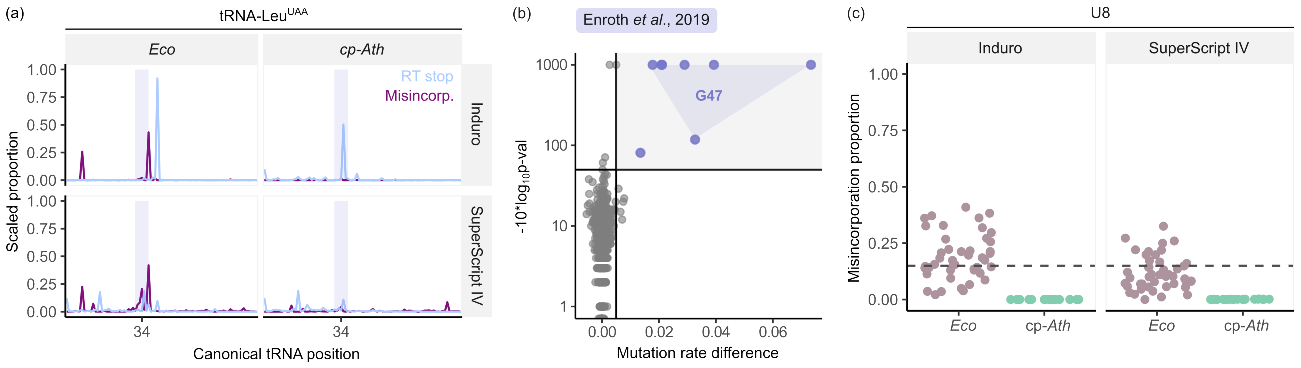


**Figure S3. Chloroplast tRNA modification landscape has similarities and distinctions from the prokaryotic system.** (**a**) Line plots showing RT misincorporations (purple) and RT stops (blue) for *E. coli* (left) and Arabidopsis chloroplasts (right) generated with both Induro (top) and SuperScript IV (bottom) RTs in tRNA-Leu$^{UAA}$. (**b**) Sodium borohydride treatment and mutational profiling were used to detect $N^7$-methylguanosine (m$^7$G) (1). Plots of mutation rate difference (x-axis) versus -10*log$_{10}$(p-val) (y-axis) for chloroplast tRNAs are presented. Most m$^7$G modifications were detected at position 47 (light blue triangle). (**c**) Dot plots depicting the proportion of RT misincorporations for position U8. Data are presented for *E. coli* (light brown) and Arabidopsis chloroplasts (green), using both Induro (left) and SuperScript IV (right) RTs. Points above the dashed line (RT misincorporation proportion $> 0.15$) likely represent modified nucleosides in these tRNAs. Notably, only *E. coli* tRNAs exhibit modification at the position.

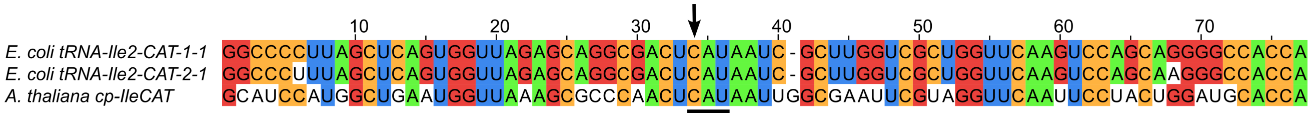


**Figure S4. Sequence context surrounding the wobble cytidine (C) in tRNA-Ile$^{CAU}$ from *E. coli* and Arabidopsis chloroplasts.** An alignment of two *E. coli* tRNAs and one Arabidopsis chloroplast tRNA is shown. The anticodon is indicated by a horizontal line, and the wobble C is marked with an arrow. Notably, the sequence flanking C34 is identical in all analysed tRNAs.

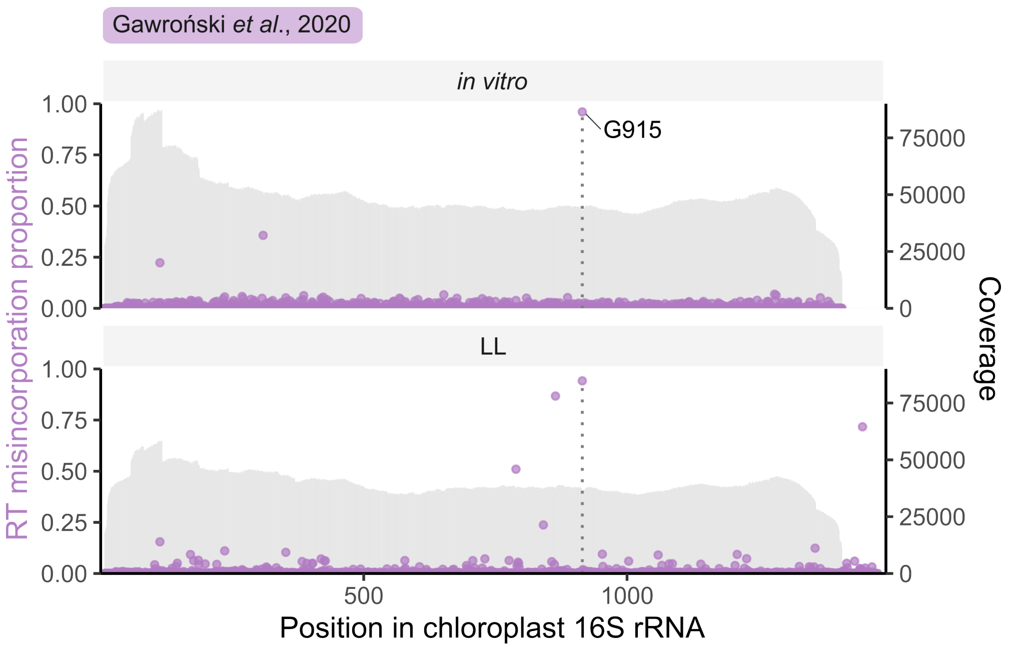


**Figure S5.** Mutational profiling was used to detect $m^2$G915 in 16S (2). The fraction of reverse transcriptase (TGIRT-III) misincorporation is shown on the left y-axis and corresponds to positions in 16S rRNA indicated on the x-axis. Read coverage is illustrated on the right y-axis. Sites with a minimum coverage of 500 reads were analysed. The results represent the average from two replicates. The *in vitro* folded and *in vivo* [grown in low light (LL) conditions] samples are presented.

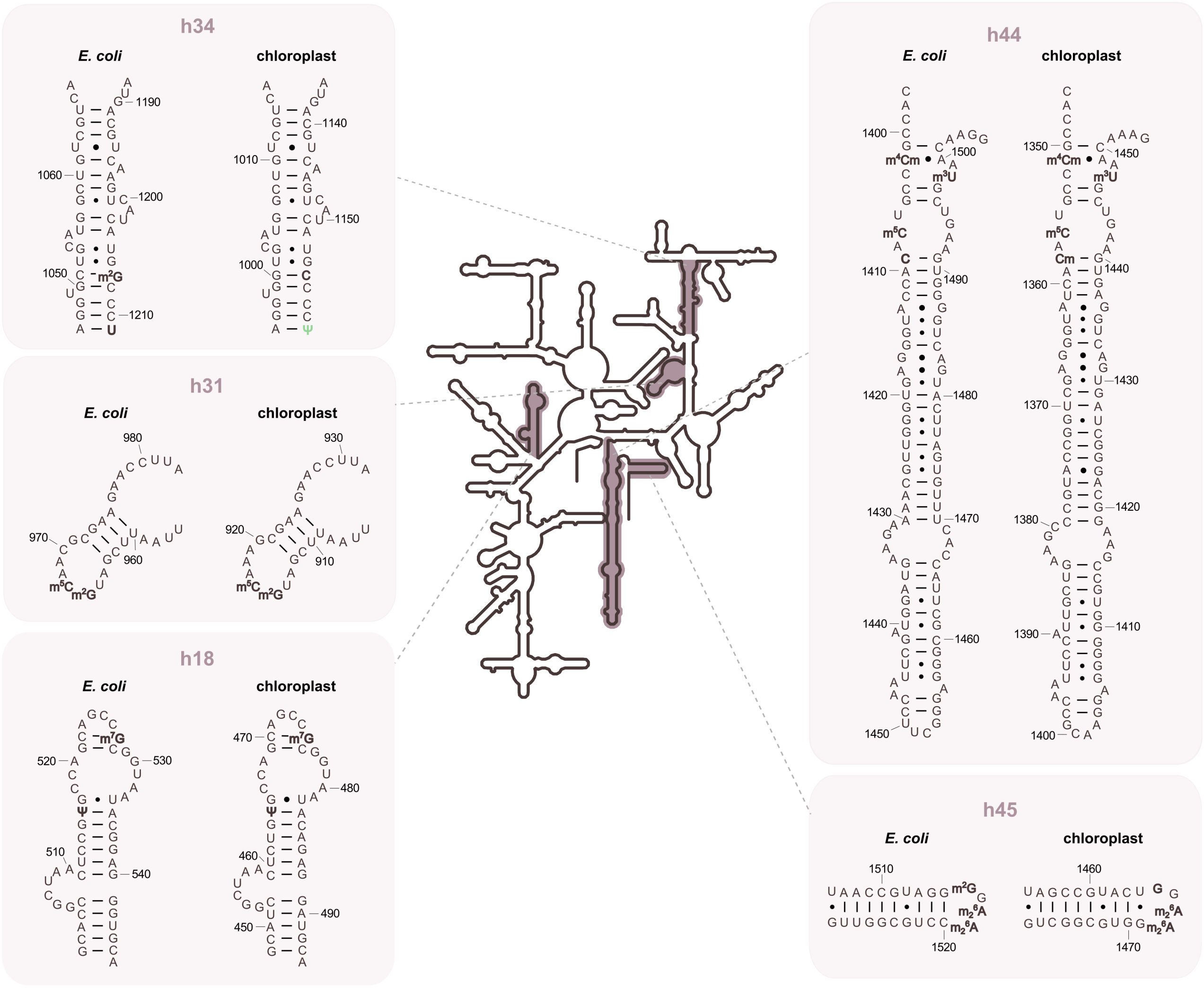


**Figure S6.** The secondary structure of *E. coli* and *A. thaliana* 16S rRNA and sequence comparison of the modified loops.

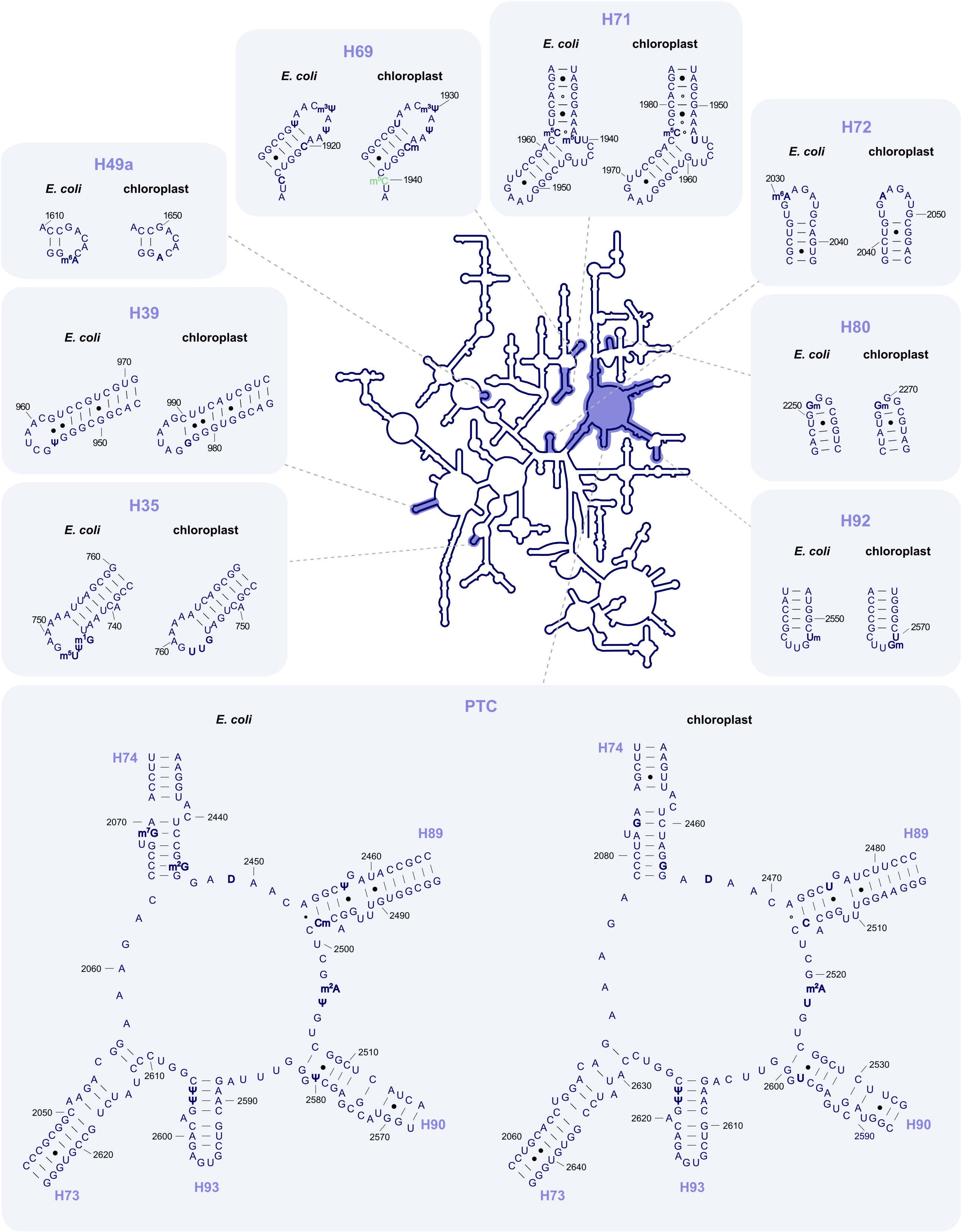


**Figure S7.** The secondary structure of *E. coli* and *A. thaliana* 23S rRNA and sequence comparison of the modified loops.

### SUPPLEMENTARY REFERENCES

(1–7)


1. Enroth,C., Poulsen,L.D., Iversen,S., Kirpekar,F., Albrechtsen,A. and Vinther,J. (2019) Detection of internal N7-methylguanosine (m7G) RNA modifications by mutational profiling sequencing. *Nucleic Acids Research*, **47**, e126–e126.
2. Gawroński,P., Pałac,A. and Scharff,L.B. (2020) Secondary Structure of Chloroplast mRNAs In Vivo and In Vitro. *Plants*, **9**, 323.
3. Sun,L., Xu,Y., Bai,S., Bai,X., Zhu,H., Dong,H., Wang,W., Zhu,X., Hao,F. and Song,C.-P. (2019) Transcriptome-wide analysis of pseudouridylation of mRNA and non-coding RNAs in Arabidopsis. *Journal of Experimental Botany*, **70**, 5089–5600.
4. Duan,H.-C., Zhang,C., Song,P., Yang,J., Wang,Y. and Jia,G. (2024) C2-methyladenosine in tRNA promotes protein translation by facilitating the decoding of tandem m2A-tRNA-dependent codons. *Nature Communications*, **15**, 1025.
5. David,R., Burgess,A., Parker,B., Li,J., Pulsford,K., Sibbritt,T., Preiss,T. and Searle,I.R. (2017) Transcriptome-Wide Mapping of RNA 5-Methylcytosine in Arabidopsis mRNAs and Noncoding RNAs. *The Plant Cell*, **29**, 445–460.
6. Burgess,A.L., David,R. and Searle,I.R. (2015) Conservation of tRNA and rRNA 5-methylcytosine in the kingdom Plantae. *BMC Plant Biology*, **15**, 199.
7. Zou,M., Mu,Y., Chai,X., Ouyang,M., Yu,L.-J., Zhang,L., Meurer,J. and Chi,W. (2020) The critical function of the plastid rRNA methyltransferase, CMAL, in ribosome biogenesis and plant development. *Nucleic Acids Research*, **48**, 3195–3210.
